# Targeting WNK1 Releases Differentiation Block in Acute Myeloid Leukemia

**DOI:** 10.64898/2026.04.22.720037

**Authors:** Jordan D. Cress, Emily M. Katoni, Parameswaran Ramakrishnan

**Affiliations:** Department of Pathology, Case Western Reserve University, 2103 Cornell Road, Cleveland, Ohio 44106, USA; The Case Comprehensive Cancer Center, Case Western Reserve University, 2103 Cornell Road, Cleveland, Ohio 44106, USA; Department of Biochemistry, Case Western Reserve University, 2109 Adelbert Road, Cleveland, Ohio 44106, USA; University Hospitals-Cleveland Medical Center, 11100 Euclid Ave, Cleveland, Ohio 44106, USA; Louis Stokes Veterans Affairs Medical Center, 10701 East Blvd, Cleveland, Ohio 44106, USA

## Abstract

Impaired differentiation is a hallmark of Acute Myeloid Leukemia (AML). Current differentiation therapies benefit only a small subset of AML patients, leaving a substantial gap in care for other subtypes. Identifying novel molecular drivers of maturation arrest is critical to expand differentiation induction to a broader range of AML patients. This study addresses this unmet clinical need, by identifying With-no-Lysine(K) kinase 1 (WNK1) as a novel regulator of AML differentiation arrest. We show that WNK1 expression and activity are elevated in AML patients. WNK1 inhibition induced differentiation accompanied by decreased growth and survival of AML cell lines and patient cells. It also inhibited self-renewal of AML patient cells in vitro and elicited significant anti-tumor activity in vivo in mouse models. Mechanistically, WNK1 inhibition derepressed the MEK-ERK-C/EBPβ signaling axis and increased the expression of myeloid differentiation genes. Our findings reveal a novel role of WNK1 in promoting AML through differentiation arrest, posing WNK1 inhibition as a potential approach for AML differentiation therapy.

## Introduction

Acute Myeloid Leukemia (AML) is the most common form of acute leukemia in adults *(1)*. It develops from the transformation of myeloid precursor cells causing hyperproliferation and differentiation arrest *^(2)^*. The current standard of care for AML treatment is chemotherapy, which targets cell proliferation and leads to off-target effects on healthy, rapidly dividing cells *(3)*. Chemotherapy is frequently ineffective at treating AML patients due to high rates of chemoresistance *(4)*. A wide array of mutations in AML leads to a high degree of heterogeneity within the disease. Many of these mutations promote chemotherapy resistance further decreasing the likelihood that patients with these mutations will respond to chemotherapy *(5)*. Due to suboptimal treatment options, the survival rate of this disease is relatively poor (5-year survival rate around 30%) *(6)*. Therefore, additional AML therapeutics are necessary to overcome the current limitations of chemotherapy.

Differentiation therapies have emerged as a promising alternative to chemotherapy. In contrast to targeting rapidly dividing cells, differentiation therapies target molecular abnormalities that block AML differentiation. The theoretical advantages of this approach include fewer off-target effects on healthy cells and overcoming drug-resistance of AML cells *^(7)^*. Releasing the differentiation block in AML counteracts other key disease features, as it has shown to decrease proliferation and induce apoptosis *^(8)^*. The success of differentiation therapy for AML treatment has been demonstrated by All-Trans Retinoic Acid (ATRA), which has displayed impressive cure rates (around 80%) in a subset of AML patients *(9)*. However, the success of this treatment has been restricted to patients with Acute promyelocytic leukemia (APL) *(10, 11)*. This highlights the need to gain a greater understanding of molecular abnormalities that control differentiation arrest in other AML subtypes to identify novel differentiation therapy targets for non-APL patients.

Kinases have been implicated in AML differentiation arrest in addition to their roles in growth and survival *(12)*. For example, FLT3 inhibitors have demonstrated the ability to induce AML differentiation as FLT3 suppresses C/EBPα activity via inhibitory phosphorylation *(13, 14)*. Another kinase, Nemo-like Kinase (NLK) also regulated myeloid differentiation by increasing C/EBPα levels. Co-targeting NLK and FLT3 further enhanced differentiation compared to NLK alone *(15)*. Additionally, Glycogen Synthase Kinase-3 (GSK3) inhibition has shown to induce AML cell differentiation and halt growth without impairing normal hematopoietic cell proliferation through activating MAPK signaling *(16)*. Therefore, targeting kinases appears to be a promising approach to induce differentiation, offering a viable therapeutic strategy for AML.

With-no-Lysine(K) kinase 1 (WNK1) has primarily been characterized for its ability to regulate ion homeostasis by controlling activity of cation-chloride cotransporters (CCCs). This process is mediated through WNK1 phosphorylation of SPAK and OSR1 which then phosphorylate CCCs, promoting ion transport across the cell membrane *(17, 18)*. WNK1 performs a variety of functions outside its role in regulating ion homeostasis including pro-tumorigenic roles such as proliferation, metastasis, and angiogenesis *(19–21)*. However, the mechanisms by which WNK1 promotes cancer progression are not well understood and vary depending on the cancer type *(22)*. Recently, it was shown that WNK1 promotes amino acid uptake and thereby supports AML growth *(23)*.

Here, we show that WNK1 expression and activity is elevated in AML patient cells compared to healthy donors and that WNK1 expression negatively correlates with survival. Inhibiting WNK1 through both genetic and pharmacologic inhibition induced differentiation of AML cells which was accompanied by increased cell cycle arrest and apoptosis. Genetic inhibition of WNK1 substrates, SPAK and OSR1, mirrored the effect of WNK1 inhibition. Colony Forming Unit (CFU) assays showed that inhibiting WNK1 reduced self-renewal capacity of primary AML patient cells. Targeting WNK1 eliminated AML patient cells with limited toxicity to healthy hematopoietic stem and progenitor cells (HSPCs). Oral administration of WNK1 inhibitor significantly reduced growth of human AML cells in xenograft mouse models. Mechanistically, WNK1 inhibition derepresses mitogen-activated protein kinase kinase (MEK) - Extracellular signal regulated Kinase (ERK) signaling, resulting in higher CCAAT/enhancer-binding protein β (C/EBPβ) levels and elevated transcription of myeloid differentiation genes.

## Results

### WNK1 expression is elevated in AML patients and negatively correlates with survival

To investigate the role of WNK1, SPAK, and OSR1 in AML, we analyzed their RNA expression in AML patients (n = 1005) and healthy donors (n = 755). We also included a small cohort of HSPCs from healthy donors (n = 13) as an additional comparison. We found significantly higher levels of WNK1 expression in AML patients when compared to healthy donor whole blood and HSPCs (Fig. 1A). SPAK showed a modest upregulation in AML patients as well as HSPCs compared to healthy whole blood (Fig. 1B). Interestingly, overall OSR1 expression was found lower both in AML patients and HSPCs than healthy whole blood, however, its level in AML samples were found higher compared to healthy HSPCs (Fig. 1C). Next, we examined whether WNK1, SPAK, and OSR1 expression levels correlated with overall survival of AML patients. We analyzed patient data from the TARGET database using SurvivalGenie *(24)* and found that high WNK1 expressors had decreased overall survival compared to low WNK1 expressors (Logrank p = 0.011, HR = 1.9) (Fig. 1D). High WNK1 expressors had a median overall survival of 22.1 months while median overall survival was not reached in low WNK1 expressors (Kaplan-Meier estimate = 64.1% survival) (Fig. 1D). SPAK expression also correlated with survival (Logrank p = 0.028, HR = 1.8) with high SPAK expressors displaying a median overall survival of 21.3 months which extends to 56.8 months for low SPAK expressors (Fig. 1E). OSR1 expression, however, did not show a correlation with overall survival as the survival curves for high and low OSR1 expressors were not significantly different (Logrank p = 0.275, HR = 0.74) (Fig. 1F).

**Fig. 1.**
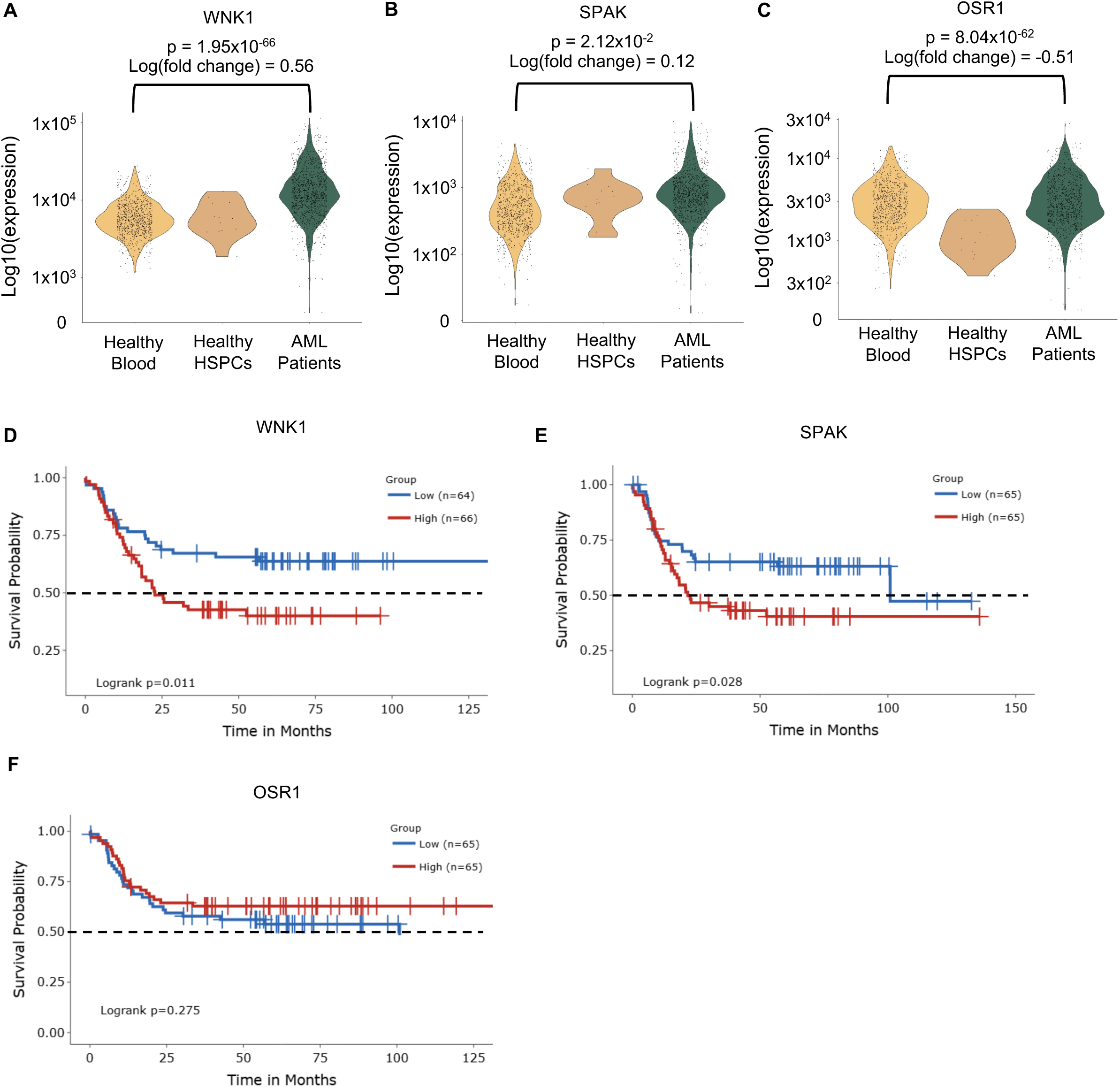
WNK1 expression is elevated in AML patients and negatively correlates with survival. A-C) Bulk RNA-sequencing comparing expression of WNK1 (A), SPAK (B), or OSR1 (C) on peripheral blood from AML patients (n=1005) and healthy donors (n = 755), as well as healthy HSPCs (n = 13). D-F) Kaplan-Meir plot showing overall survival of AML patients from the TARGET dataset with high (top 20%) (n = 66) or low (bottom 20%) (n = 64) WNK1 (D), SPAK (E), and OSR1 (F) expression.

### WNK1, SPAK, and OSR1 are involved in the regulation of cell cycle arrest, apoptosis, and differentiation of AML cells

Since we observed elevated activation of WNK1, SPAK, and OSR1 in AML, we next aimed to explore the potential pro-leukemic effects of these proteins. To do this, we genetically suppressed WNK1, SPAK (gene name: STK39), and OSR1 (gene name: OXSR1) in MOLM-14 cells. Knocking down WNK1 ablated the phospho-SPAK/OSR1 signal while knocking down SPAK did not affect the phosphorylation of OSR1 and vice-versa (Fig. 2A). Interestingly, knocking down WNK1, SPAK, or OSR1 reduced AML proliferation over the course of 4 days to similar levels (Fig. 2B). Cell cycle analysis revealed that knocking down WNK1, SPAK, or OSR1 resulted in an increased arrest of cells in G1 phase (Fig. 2C). WNK1, SPAK, and OSR1 knockdown also induced apoptosis of MOLM-14 cells. We observed that SPAK deficiency caused slightly higher levels of apoptosis than WNK1 or OSR1 deficiencies (Fig. 2D).

**Fig. 2.**
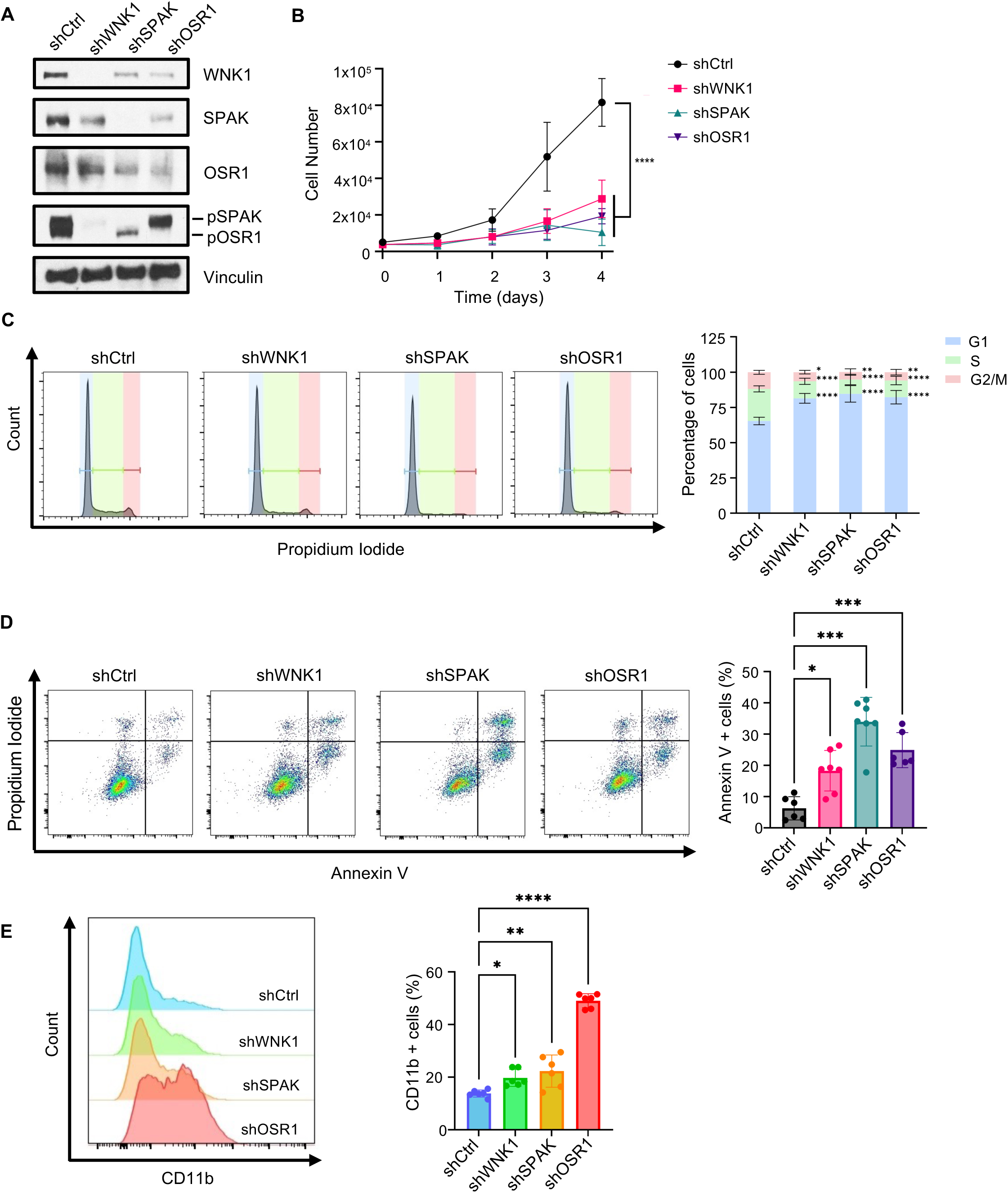
WNK1, SPAK, and OSR1 knockdown induces cell cycle arrest, apoptosis, and differentiation of AML cells. A) Western blot validating decreased protein levels of WNK1, pSPAK/OSR1 (S373/S325), SPAK, and OSR1 after lentiviral shRNA knockdown of MOLM-14 cells. B) Growth of shCtrl, shWNK1, shSPAK, and shOSR1 MOLM-14 cells over the course of 4 days quantified by automated cell counting. C-E) Cells were assessed 4 days after lentiviral transduction. C) Cell cycle analysis (Left) Histogram showing cellular DNA content by propidium iodide staining., (Right) Percentages of cells in each phase of cell cycle. D) (Left) Annexin V and PI staining for apoptosis, (Right) Percentage of Annexin V positive cells. E) (Left) Histogram of CD11b surface expression, (Right) Percentage of CD11b+ cells. Statistical analysis was done using a one-way ANOVA (B, D-E) or a two-way ANOVA (C) with Dunnet’s multiple comparisons test. *, P < 0.05; **, P < 0.01; ***, P < 0.001; ****, P < 0.0001.

We also studied the role of WNK1, SPAK, and OSR1 in AML differentiation arrest by examining whether knocking down these proteins induced differentiation. Knocking down WNK1, SPAK, or OSR1 caused increased differentiation of MOLM-14 cells as indicated by enhanced CD11b expression (Fig. 2E). This differentiation appears to be primarily granulocytic as only a modest increase was observed in CD14 expression (fig. S1A-). Interestingly, OSR1 knockdown showed much higher CD11b positivity than WNK1 and SPAK knockdowns.

### Pharmacological targeting of WNK1 induces cell cycle arrest, apoptosis, and differentiation of AML cells

To determine whether pharmacological targeting of WNK1 phenocopies the effects of its genetic depletion, and whether its kinase activity is involved in leukemogenic functions, we utilized WNK-IN-11, an allosteric kinase inhibitor optimized for WNK1 specificity *(25)*. WNK-IN-11 inhibited the constitutive phosphorylation of SPAK and OSR1 in MOLM-14 and OCI-AML3 cells (Fig. 3A-B). Next, we examined the dose-dependent effect of WNK-IN-11 on cell viability and found that OCI-AML3 cells (IC_50_ = 4.85 µM) were more sensitive to WNK-IN-11 compared to MOLM-14 cells (IC_50_ = 6.09 µM) (Fig. 3C). Maximum inhibition was observed at 10 µM (Fig. 3C) and this concentration was used for all subsequent experiments. Our kinetic analysis showed that WNK-IN-11 efficiently suppressed phosphorylation of SPAK/OSR1 up to 24 hours which started to reappear by 48 hours post-treatment (Fig. 3D). Based on this kinetic study, to maintain sustained WNK1-SPAK/OSR1 inhibition, studies longer than 24 hours were supplemented with WNK-IN-11 daily. We found no increase in the numbers of MOLM-14 and OCI-AML cells treated with WNK-IN-11 over the course of 4 days indicating significant growth inhibition (Fig. 3E-F). WNK-IN-11 also disrupted cell cycle progression through inducing G1 arrest in MOLM-14 and OCI-AML3 cells (Fig. 3G-H) as well as THP-1 and MOLM-13 cell lines (fig. S1B-C).

**Fig. 3.**
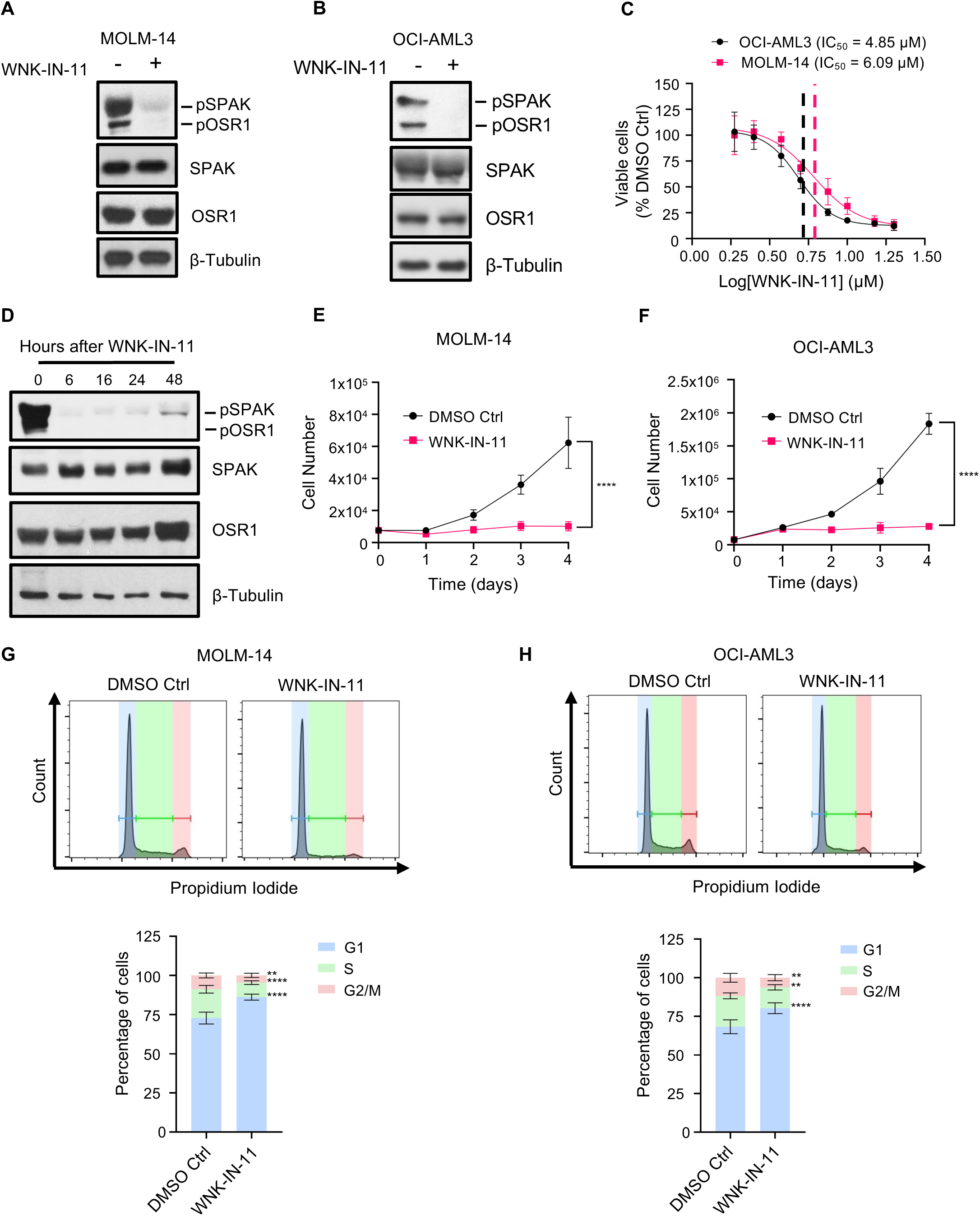
WNK1 inhibition reduces AML proliferation and promotes cell cycle arrest. A-B) MOLM-14 (A) and OCI-AML3 (B) cells were treated with WNK-IN-11 (10 µM) or DMSO control for 3 hours and WNK1 activity was measured through phosphorylation of SPAK and OSR1 (S373/S325). C) Dose-response curve showing the effect of increasing WNK-IN-11 concentrations on MOLM-14 and OCI-AML3 cell viability after 4 days of treatment. D) Phosphorylation of SPAK and OSR1 (S373/S325) assessed from 6-48 hours after WNK-IN-11 treatment in OCI-AML3 cells. E-F) MOLM-14 (E) and OCI-AML3 (F) cells were treated with WNK-IN-11 (10 µM) or DMSO control and viable cell counts over the course of 4 days were quantified by automated cell counting. G-H) MOLM-14 (G) and OCI-AML3 (H) cells were treated with WNK-IN-11 (10 µM) or DMSO control for 3 days prior to cell cycle analysis, (Top) Histogram showing cellular DNA content by propidium iodide staining. (Bottom) Percentages of cells in each phase of cell cycle. Statistical analysis was done using a Student’s t-test (E-F) or a two-way ANOVA with Šídák’s multiple comparisons test (G-H). **, P < 0.01, ****, P < 0.0001.

When flow cytometry-based cell cycle analysis was performed on total events (not excluding dead cells) WNK-IN-11 produced a prominent population of cells in the G0 phase indicating significant DNA fragmentation and apoptosis (fig. S1D-G). To support that apoptosis was occurring, we also saw that WNK-IN-11 increased Annexin V positivity in AML cell lines (Fig. 4A-B, fig. S2A-B). Interestingly, the OCI-AML3 and MOLM-13 cells were more susceptible to WNK-IN-11 induced apoptosis than MOLM-14 and THP-1 cells as indicated by higher Annexin V positivity (Fig. 4A-B, fig. S2A-B). We investigated if the resistance to WNK-IN-11 induced apoptosis was due to higher levels of WNK1 and found that while MOLM-14 cells had higher WNK1 levels, THP-1 cells had equivalent WNK1 levels with the more WNK-IN-11 susceptible cell lines (OCI-AML3 and MOLM-13) suggesting that differential WNK1 levels cannot completely explain these effects alone (fig. S2C). However, doubling the concentration of WNK-IN-11 to 20 µM led to an increase in Annexin V positive staining of MOLM-14 cells similar to the level of OCI-AML3 cells treated with 10 µM WNK-IN-11 (fig. S2D-E). WNK-IN-11 treatment induced PARP and Caspase 3 cleavage (Fig. 4C-D) and pan-caspase inhibition with Z-VAD partially blocked apoptosis further demonstrating caspase cascade activation and apoptotic cell death (fig. S2F).

**Fig. 4.**
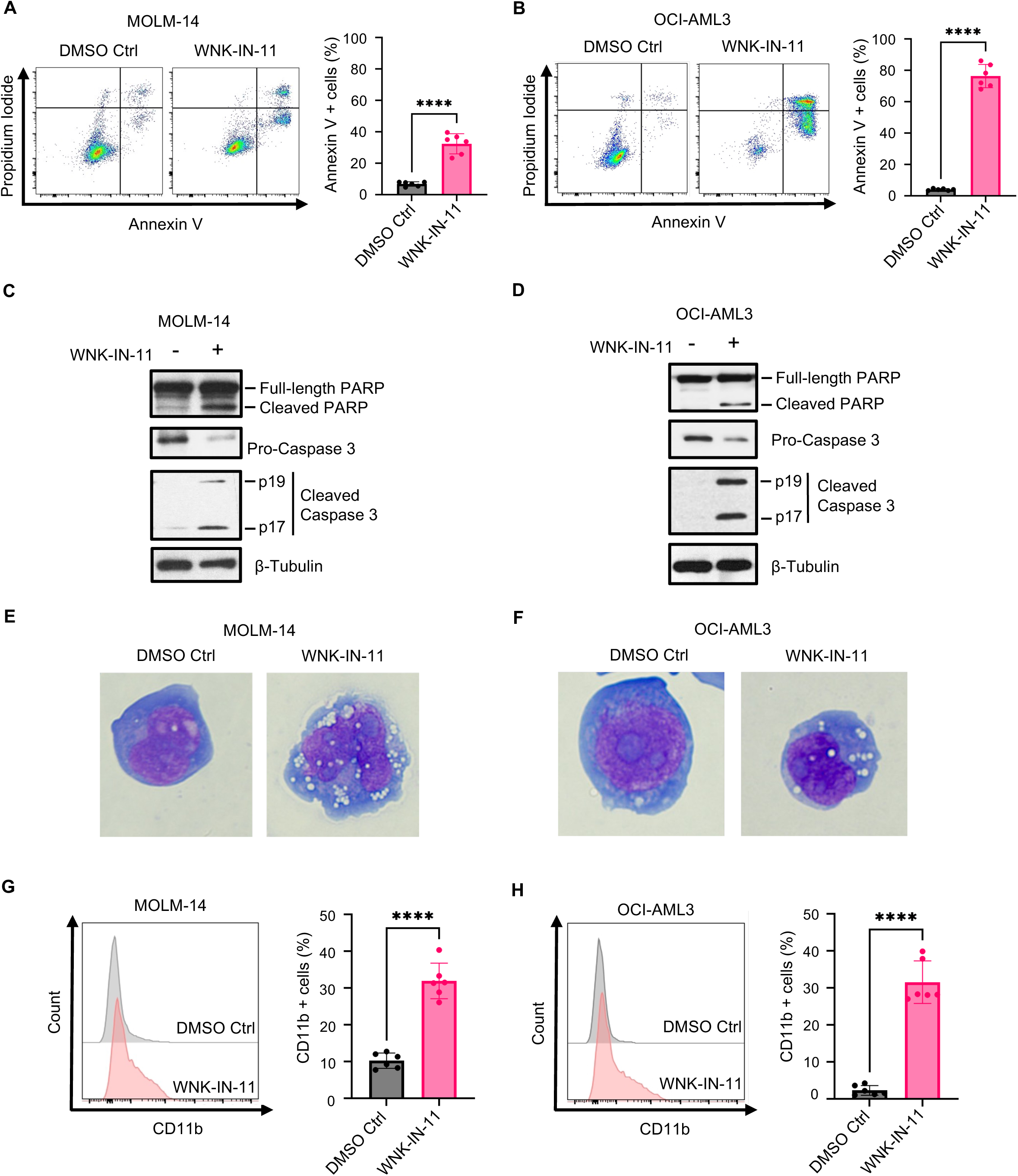
WNK1 inhibition induces AML apoptosis and differentiation. A-B) Apoptosis markers were assessed on MOLM-14 (A) and OCI-AML3 (B) cells after 4 days of WNK-IN-11 (10 µM) or DMSO Control treatment. (Left) Annexin V and PI staining for apoptosis. (Right) Percentage of Annexin V positive cells. C-D) Western blot showing PARP and Caspase 3 cleavage after 3 days of WNK-IN-11 (10 µM) treatment in MOLM-14 (C) and OCI-AML3 (D) cells. E-F) Wright-Giemsa stain showing morphological changes of MOLM-14 (E) and OCI-AML3 (F) 3 days after treatment with WNK-IN-11 (10 µM) or DMSO control (magnification: 100x). G-H) CD11b surface expression of MOLM-14 (G) and OCI-AML3 (H) cells 3 days after treatment with WNK-IN-11 (10 µM) or DMSO control. (Left) Histogram of CD11b surface expression, (Right) Percentage of CD11b+ cells. Statistical analysis was done using a Student’s t-test. ****, P < 0.0001.

Next, we assessed the effect of WNK1 inhibition on differentiation of AML cells. We assessed morphological changes of MOLM-14 and OCI-AML3 cells after WNK1 inhibition and observed changes such as nuclear indentation, nuclear condensation, and increased vacuolization (Fig. 4E-F, fig. S2G-H), all of which are defining features of myeloid differentiation *(26, 27)*. Further characterization of the differentiated cells showed that inhibiting WNK1 primarily produced CD11b+CD14- cells (∼30%) and a small number of CD11b^+^CD14^+^ cells (∼5%) suggesting predominant granulocytic differentiation over monocytic differentiation (Fig. 4G-H, fig. S2I-J). This predominant CD11b^+^CD14^-^ population after WNK1 inhibition was also observed in THP-1 and MOLM-13 cells with THP-1 cells showing slightly higher levels of CD11b^+^CD14^+^ cells (∼15%) (fig. S3A-B).

### WNK1 inhibition reduces leukemic burden in mice

To evaluate the in vivo anti-leukemic effect of WNK1 inhibition, we performed mouse xenograft studies. Because WNK-IN-11 has previously shown to have poor bioavailability*(25)*, we used the pan-WNK inhibitor, WNK-463 *(28)* as an alternative. First, we tested toxicity of WNK-463 treatment and found that it did not significantly alter mouse body weight over the course of a 3-week treatment period (fig. S3C). For the xenograft assay, luciferase-expressing MOLM-14 and OCI-AML3 cells were transplanted into NOD-scid IL2Rg^null^ (NSG) mice via tail vein injection. After 7 days, engraftment was confirmed using bioluminescent imaging and mice were treated for 2 weeks with WNK-463 (1.5 mg/kg) or Vehicle Control. Bioluminescent images were taken at days 7, 14, and 21 post-injection (Fig. 5A). WNK1 inhibition significantly reduced leukemic burden in the mice as evidenced by reduced bioluminescent signal compared to the vehicle control (Fig. 5B-E).

**Fig. 5.**
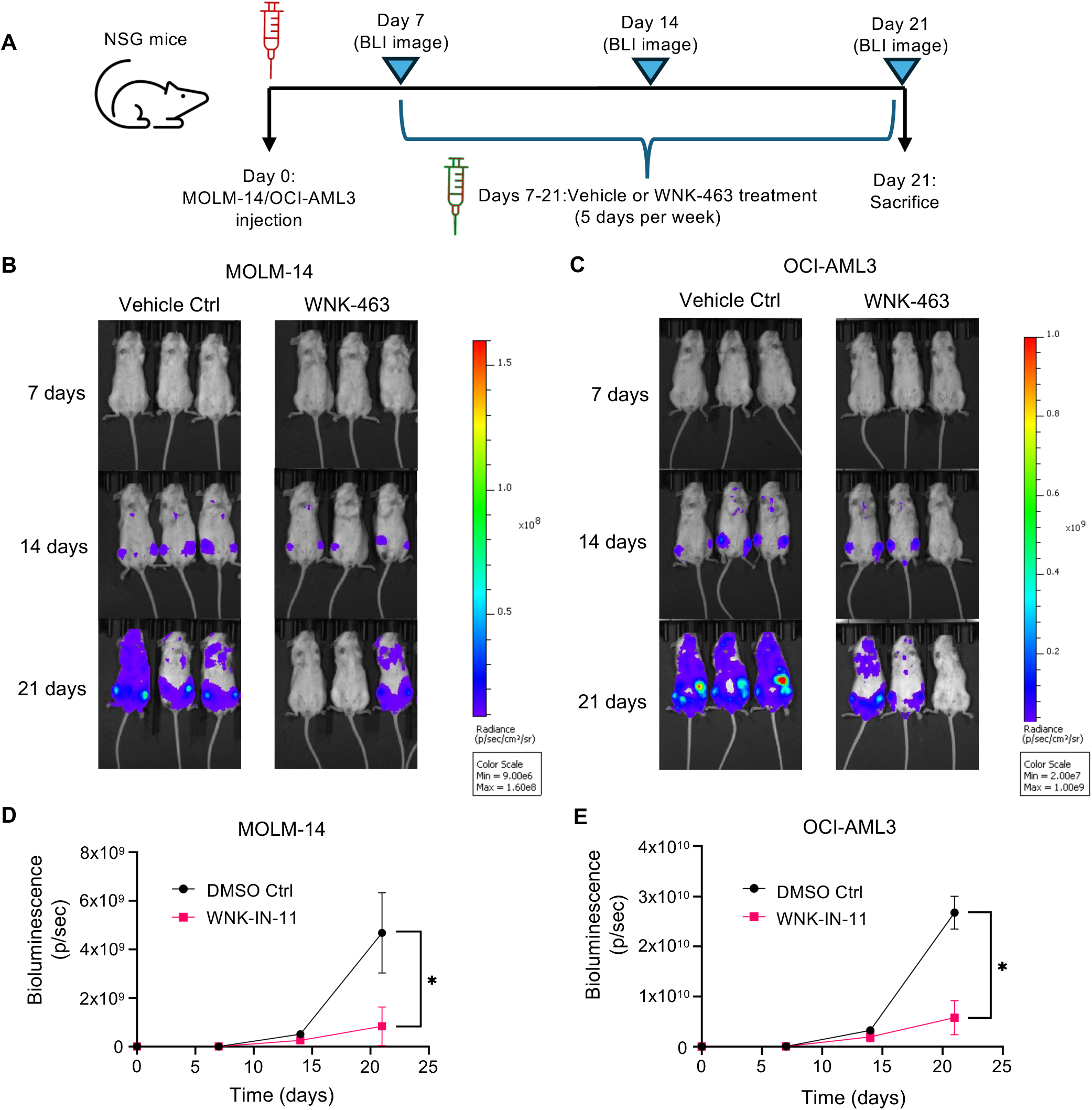
WNK1 inhibition reduces leukemic burden in mice and induces death and differentiation of primary AML cells. A) Overview of experiment design for xenograft study. NSG mice were intravenously injected with MOLM-14 or OCI-AML3 cells. After 7 days, mice were administered WNK-463 (1.5 mg/kg) or vehicle control via oral gavage 5 days a week for two weeks. B) Bioluminescent images of mice at days 7, 14, and 21 post-injection of MOLM-14 (B) and OCI-AML3 (C) cells. D-E) Luciferase intensity plotted at each imaging timepoint for MOLM-14 (D) and OCI-AML3 (E) xenografts. Statistical analysis was done using a Student’s t-test on day 21 data (D-E) *, P < 0.05.

### WNK1 inhibition impairs self-renewal capacity and induces differentiation and death of patient-derived AML cells

To translate our AML cell line findings into clinically relevant evidence, we further investigated the ability of WNK-IN-11 to restrict AML progression using patient samples (for patient demographics, see Table S1). We examined WNK1, SPAK, and OSR1 protein expression in AML patient cells and healthy HSPCs. We found that AML cells expressed significantly higher WNK1 protein (Fig. 6A), substantiating our transcriptome analysis (Fig. 1A). SPAK was slightly elevated in AML samples, while OSR1 levels were comparable in AML cells and healthy HSPCs (Fig. 6A). Notably, we found higher phosphorylation of SPAK and OSR1 in AML, suggesting increased WNK1 kinase activity to correlate with its elevated expression (Fig. 6A).

**Fig. 6.**
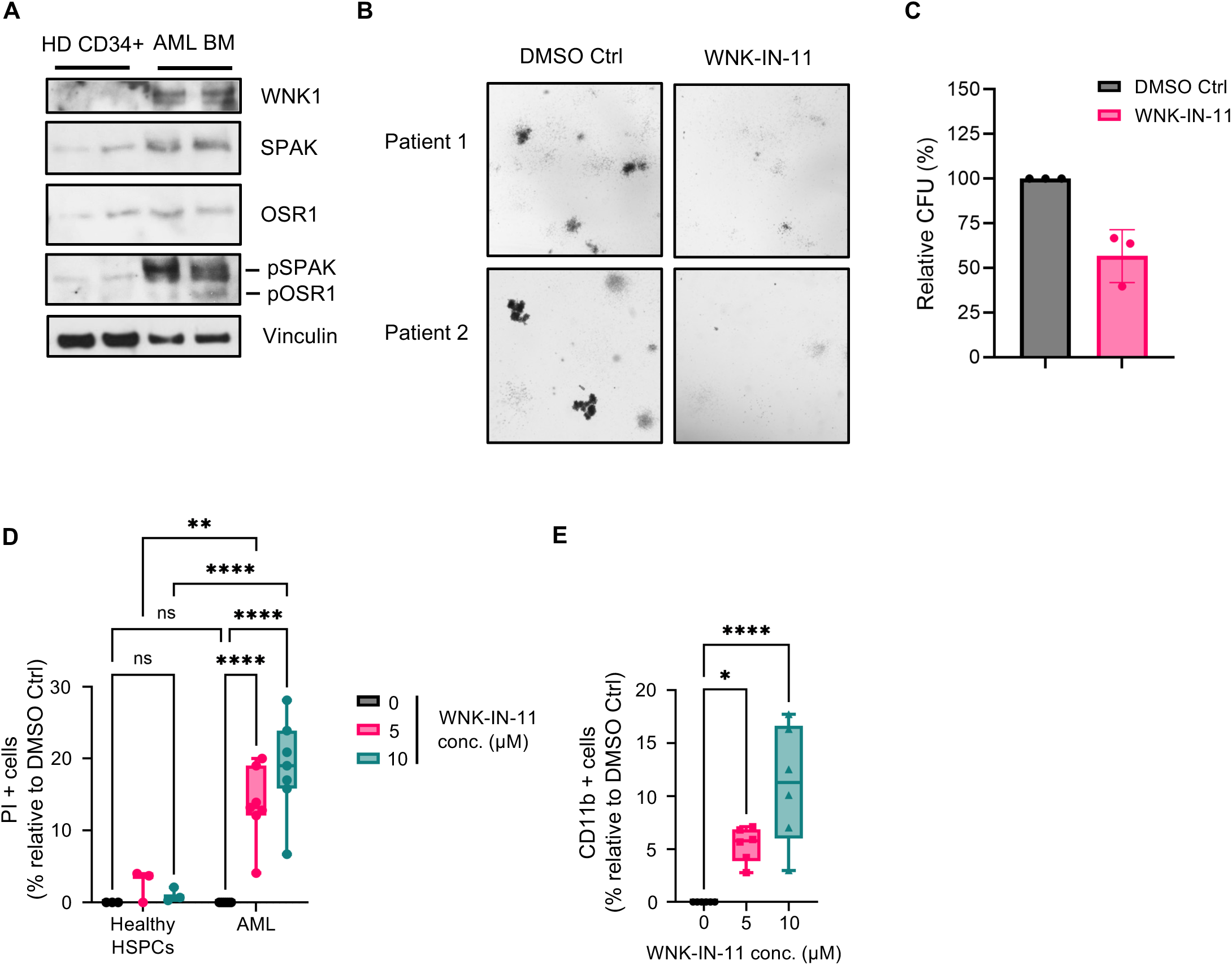
Targeting WNK1 induces cell cycle arrest, death, and differentiation of AML patient cells while sparing healthy HSPCs. A) Western blot comparing WNK1, pSPAK/OSR1 (S373/S325), SPAK, and OSR1 levels in CD34+ HSPCs from healthy patients and AML bone marrow samples. B-C) Representative images (B) and relative quantification (C) of AML MNC colony formation treated with WNK-IN-11 (10 µM) or DMSO control 14 days after seeding in MethoCult media. D) Cell death of healthy HSPCs (n=3) and AML MNCs (n=7) after 4 days of WNK-IN-11 (5 and 10 µM) treatment normalized to the DMSO control. Technical duplicates from each donor were averaged and plotted as a single data point. E) CD11b expression of AML MNCs (n=3) 3 days after 10 µM WNK-IN-11 treatment normalized to DMSO control. Statistical analysis was done using a Student’s t-test (C), a two-way ANOVA with Šídák’s multiple comparisons test (D), or one-way ANOVA with Dunnet’s multiple comparisons test (E). *, P < 0.05; **, P < 0.01; ****, P < 0.0001; ns, not significant.

Next, we evaluated the impact of WNK1 inhibition on clonal proliferation using bone marrow cells derived from AML patients in a colony forming unit (CFU) assay. We found that WNK-IN-11 treatment reduced AML self-renewal capacity as evidenced by significantly reduced colony formation (Fig. 6B-C). To assess the ability of WNK inhibition to specifically eliminate AML cells, we compared the effects of WNK-IN-11 treatment on CD34^+^ HSPCs purified from healthy individuals (fig. S3D) and AML patient cells. We found that WNK-IN-11 induced death in AML cells with limited toxicity towards healthy HSPCs (Fig. 6D). WNK-IN-11 treatment also induced differentiation of AML blasts as indicated by increased CD11b positivity (Fig. 6E).

### WNK1 inhibition induces AML differentiation by increasing C/EBPβ expression and activity

The canonical WNK1 signaling pathway (WNK1-SPAK/OSR1-NKCC1/NKCC2) does not account for its pro-leukemic functions as neither knockout of NKCC1 nor NKCC2 results in decreased survival of AML or other cancers *(23)*. We also found that pharmacological targeting of NKCC1 and NKCC2 with Bumetanide inhibited chloride influx (fig. S3E), but showed no effect on AML survival, cell cycle regulation, or differentiation (fig. S3F-H). This posits the existence of non-canonical routes downstream of WNK1 that potentially govern its pro-leukemic functions. To investigate this, we employed RNA sequencing to identify gene expression changes in OCI-AML3 cells treated with WNK-IN-11. Differential gene expression analysis revealed that expression of 977 genes was altered after WNK1 inhibition, 777 of which were upregulated and 200 that were downregulated (Fig. 7A). The differential expression profile observed between the upregulated and downregulated genes suggests that the effect of WNK1 is predominantly repressive, exhibiting about 4-fold greater suppressor activity than its ability to positively regulate gene expression. We performed KEGG pathway enrichment analysis to categorize the genes upregulated upon WNK-IN-11 treatment into functional pathways. We found substantial enrichment of genes involved in immune related functions such as cytokine interactions, complement cascade response, and cell adhesion (Fig. 7B). We also observed hematopoietic cell lineage gene set enrichment, which included multiple myeloid differentiation markers such as CSF1R, CD38, ITGAM (CD11b), and CD14 (Fig. 7B). This suggested plausible WNK1-dependent suppression of transcription factors controlling myeloid differentiation. To explore this, we further screened the expression of a panel of myeloid differentiation associated transcription factors identified in our RNA sequencing dataset. While most of the transcription factors were not significantly altered following WNK1 inhibition, we observed a significant increase in C/EBPβ expression (Fig. 7C). C/EBPβ has been shown to be prominently upregulated in response to other differentiation therapies such as ATRA and 1α,25-dihydroxyvitamin D (1,25D) and promotes expression of differentiation markers such as CD11b *^(29, 30)^*. To investigate whether increased C/EBPβ expression correlated with increased C/EBPβ activity, we performed a gene set enrichment analysis on C/EBPβ transcriptional targets. WNK1 inhibition led to a significant enrichment of C/EBPβ target genes compared to the control (q = 0.2118) (fig. S4A). We validated the increase in C/EBPβ expression through RT-qPCR and found that WNK1 inhibition induced a 4-fold enrichment (Fig. 7D). This increased expression was also reflected on the protein level where we saw an increase in both transcriptionally active C/EBPβ isoforms (Liver-enriched Activating Protein 1/2; also called LAP and LAP*) (Fig. 7E).

**Fig. 7.**
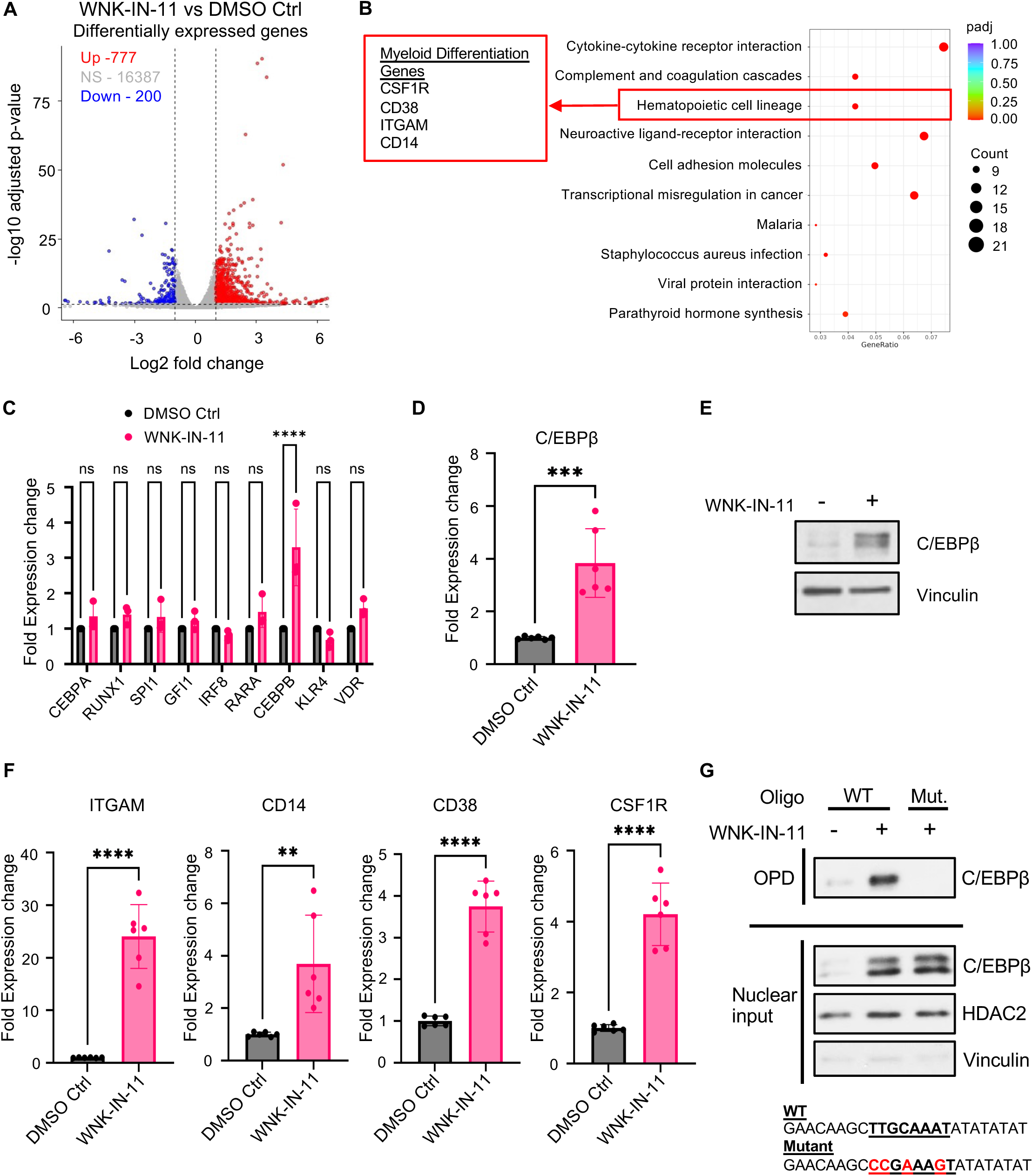
WNK1 inhibition induces AML differentiation by increasing C/EBPβ activity and expression. A) Volcano plot showing differential gene expression of OCI-AML3 cells treated with 10 µM WNK-IN-11 for 12 hours compared to the DMSO control. B) KEGG pathway analysis showing the top 10 differentially regulated pathways in WNK-IN-11 treatment compared to DMSO control. C) Expression of indicated transcription factors known to regulate myeloid differentiation after 12 hours of 10 µM WNK-IN-11 treatment. D) C/EBPβ mRNA expression of OCI-AML3 cells treated with WNK-IN-11 (10 µM) or DMSO control for 24 hours. E) C/EBPβ protein expression of OCI-AML3 cells treated with WNK-IN-11 (10 µM) or DMSO control for 24 hours. F) mRNA expression of myeloid differentiation genes, ITGAM, CD14, CD38 and CSF1R, induced after 24 hours of WNK-IN-11 treatment (10 µM) analyzed via qPCR. G) Oligonucleotide pull-down showing C/EBPβ DNA binding to a C/EBPβ binding motif in the CD11b (gene name: ITGAM) promoter region (TTGCAAAT) in OCI-AML3 cells 6 hours after treatment with WNK-IN-11 (10 µM) or DMSO control. Statistical analysis was done using a two-way ANOVA with Šídák’s multiple comparisons test (C) or a Student’s t-test (D, F). **, P < 0.01; ***, P < 0.001; ****, P < 0.0001; ns, not significant.

We then used RT-qPCR to validate expression levels of the myeloid differentiation markers we identified in our RNA-sequencing and verified increases in ITGAM (24-fold), CD14 (4-fold), CD38 (4-fold), and CSF1R (4-fold) following WNK-IN-11 treatment (Fig. 7F). We found that the increase in CD11b (ITGAM) gene expression began to increase at 6 hours after WNK-IN-11 (fig. S4B) which preceded the increase in C/EBPβ protein expression at 24 hours (fig. S4C). This led us to hypothesize that C/EBPβ transcriptional activity was elevated prior to its global upregulation to account for this gene expression kinetic pattern. To explore this possibility, we performed an oligonucleotide pull-down experiment with biotinylated oligonucleotides containing a C/EBPβ binding motif found in the ITGAM promoter region (TTGCAAAT) (fig. S4D) to evaluate C/EBPβ DNA binding capacity. Using nuclear extracts from cells treated with WNK-IN-11 or vehicle control cells, we showed increased oligonucleotide binding of C/EBPβ following WNK1 inhibition along with increased nuclear levels of C/EBPβ (Fig. 7G). We confirmed the specificity of this C/EBPβ-DNA interaction using an oligonucleotide with mutations introduced at highly conserved sites (Fig. 7G).

### WNK1-mediated C/EBPβ suppression is regulated through MEK-ERK signaling

To identify WNK1 inhibition-dependent differentially phosphorylated proteins and signal perturbations, we employed a phosphoproteomic mass spectrometry screen on cells treated with or without WNK-IN-11 (Data file S1). From this assay, we identified 142 peptides that were differentially phosphorylated by WNK1 inhibition: 65 of which showed increased and 77 showed decreased phosphorylation (Fig. 8A). To explore which kinases had altered activation with WNK1 inhibition, we performed a kinase-substrate enrichment analysis using the KSEA App online tool *(31–34)*. The phosphoproteomics data was used to score the activity level of each kinase based on the phosphorylation status of its known and predicted substrates. This analysis revealed that WNK1 inhibition enhanced activity of multiple kinases involved in ERK signaling including RAF proto-oncogene serine/threonine-protein kinase (gene name: Raf1), MEK1 (gene name: MAP2K1), MEK2 (gene name: MAP2K2) as well as ERK2 (gene name: MAPK1) itself (Fig. 8B, Data file S2).

**Fig. 8.**
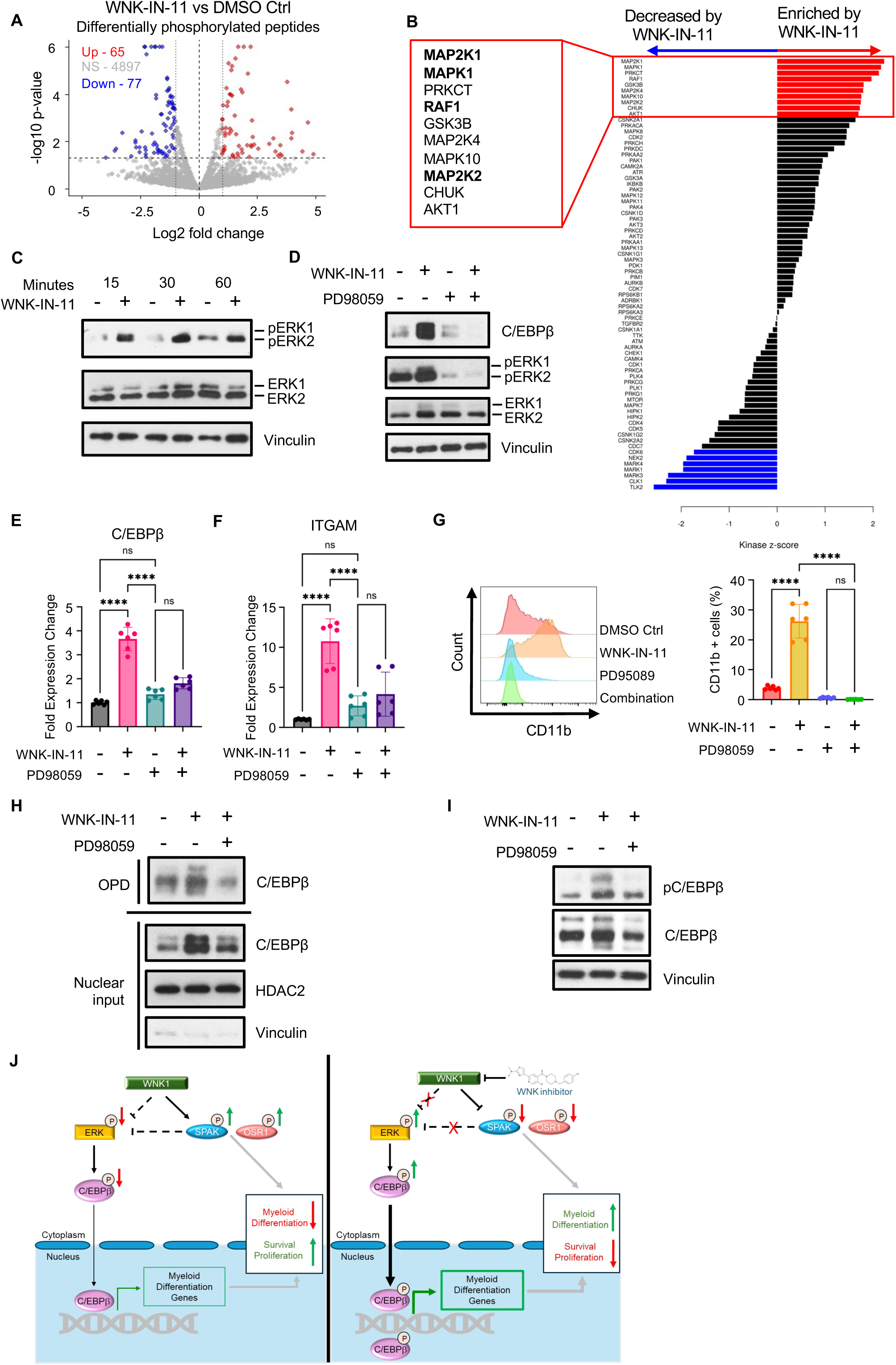
WNK1 inhibition increases ERK activation that regulates AML differentiation. A) Volcano plot showing differentially phosphorylated peptides in OCI-AML3 cells after 3 hours of treatment with WNK-IN-11 (10 µM) compared to the vehicle control. B) Kinase-substrate enrichment analysis showing upregulated (red) and downregulated (blue) kinases after WNK-IN-11 treatment based on the phosphorylation status of their known and predicted substrates. C) ERK1/2 phosphorylation in OCI-AML3 cells at T202/Y204 from 15, 30 and 60 minutes after treatment with WNK-IN-11 (10 µM) or DMSO control. D) C/EBPβ expression in OCI-AML3 cells 24 hours after treatment with WNK-IN-11 (10 µM), PD98059 (20 µM), or a combination of both. E-F) mRNA expression of C/EBPβ (E) and ITGAM (F) in OCI-AML3 cells 24 hours after treatment with WNK-IN-11 (10 µM), PD98059 (20 µM), or a combination of both. G) CD11b surface expression on OCI-AML3 cells 24 hours after treatment with WNK-IN-11 (10 µM), PD98059 (20 µM), or a combination of both. (Left) Histogram of CD11b surface expression, (Right) Percentage of CD11b+ cells. H) Oligonucleotide pull-down showing C/EBPβ DNA binding to the C/EBPβ binding motif in the ITGAM promoter region (TTGCAAAT) in OCI-AML3 cells 6 hours after treatment with WNK-IN-11 (10 µM), PD98059 (20 µM), or a combination of both. I) C/EBPβ phosphorylation at T235 in OCI-AML3 cells 1 hour after treatment WNK-IN-11 (10 µM) alone or in combination with PD98059 (20 µM). J) Schematic model showing mechanistic role of WNK1 in AML differentiation. (Left) In AML cells, WNK1 phosphorylates SPAK and OSR1 which promotes AML survival, proliferation, and differentiation arrest. WNK1 represses ERK phosphorylation and activation (which may be dependent or independent of SPAK/OSR1). This leads to decreases in phospho-C/EBPβ, nuclear C/EBPβ localization (indicated by the thin black arrow), and C/EBPβ DNA binding, suppressing myeloid differentiation gene expression (indicated by the thin green arrow). Dashed lines represent potential direct or indirect interactions. (Right) Inhibiting WNK1 in AML cells prevents phosphorylation of SPAK and OSR1. This leads to increased differentiation and decreased survival and proliferation of AML cells. Inhibiting WNK1 derepresses ERK leading to its increased activity and increases in C/EBPβ phosphorylation, nuclear C/EBPβ localization (indicated by the thick black arrow), C/EBPβ DNA binding, and transcription of myeloid differentiation genes (indicated by the thick green arrow). Statistical analysis was done using a one-way ANOVA with Tukey’s multiple comparisons test ****, P < 0.0001; ns, not significant.

To delineate WNK1 inhibition dependent ERK phosphorylation, we examined multiple timepoints within 1 hour after adding WNK-IN-11. We observed increased phosphorylation of ERK at T202/Y204 starting at 15 minutes post-treatment and peaking at 30 minutes (Fig. 8C). Interestingly, the WNK-IN-11 predominately phosphorylated ERK2 as opposed to ERK1 which could explain why we only observed enrichment of ERK2 substrates and not ERK1 (Fig. 8B-C). As increases in ERK activity has previously been associated with AML differentiation *(16, 35)*, we investigated whether inhibiting ERK would prevent WNK-IN-11 induced differentiation through regulating C/EBPβ. Indeed, we found that inhibiting the MEK-ERK signaling with the MEK inhibitor PD98059, ablated both the protein (Fig. 8D) and RNA (Fig. 8E) level increases in C/EBPβ expression. MEK inhibition also counteracted WNK-IN-11 induced differentiation as measured by reduction of CD11b transcript (Fig. 8F) and cell surface expression (Fig. 8G). We further examined the direct role of C/EBPβ in CD11b expression through an oligonucleotide-pull down assay using the ITGAM promoter C/EBPβ binding motif (fig. S4D). We observed an increase in nuclear levels and binding of C/EBPβ to the oligonucleotide when WNK1 was inhibited, which was abolished after addition of PD98059 (Fig. 8H). Previous studies have found that ERK phosphorylates C/EBPβ at T235, enhancing its DNA binding and transcriptional activation *(36–38)*. To test whether MEK-ERK was inducing C/EBPβ phosphorylation after WNK1 inhibition, we treated cells with WNK-IN-11 alone or a combination of WNK-IN-11 and PD98059 and assessed C/EBPβ phosphorylation. We found that WNK-IN-11 induced C/EBPβ phosphorylation which was ablated by PD98059 (Fig. 8I). This shows a novel role for WNK1 as a regulator of MEK-ERK-C/EBPβ signaling to control AML differentiation (Schematic model Fig 8J).

## Discussion

In this study, we reveal WNK1 as a novel regulator of AML differentiation arrest. Genetic and pharmacological targeting of WNK1 induced primarily granulocytic differentiation like what has been observed with other differentiation inducing agents *^(8, 39)^*. We also observed that differentiation was accompanied by cell cycle arrest and apoptosis induction. WNK1 inhibition induced cell cycle arrest in G1 phase suggesting a role in regulating cell cycle checkpoints responsible for the G1-S phase transition. Our pre-clinical studies show the potential for WNK1 inhibition to target AML cells while sparing normal cells as we observed limited toxicity in healthy donor HSPCs as well as in our mouse model. Our study emphasizes the pro-leukemic role of WNK1 in AML and opens the door for its potential as a target to induce AML cell differentiation, and thereby growth arrest, apoptosis and eventual elimination.

Releasing differentiation arrest has shown promise for AML treatment. In addition to reprogramming AML cells from their malignant state, this approach can also restore immune cell populations that have been compromised *^(39)^*. At the same time, this should decrease the amounts of inflammatory cell death that typically occur with other cytotoxic agents that can lead to organ-damaging side effects like acute kidney injury *(40)*. We speculate that inducing differentiation through WNK1 inhibition will also target LSCs, which are resistant to chemotherapy due to their quiescent state *(41)*. Inducing LSC differentiation may reduce their self-renewing capacity as well as make them more chemosensitive, aiding in their clearance. Our CFU assay showed that WNK1 inhibition decreased colony formation of AML patient cells which likely indicates the ability of WNK1 inhibitors to target LSCs. However, further testing with serial replating assays and serial transplantation assays are needed to better characterize this finding. While differentiation therapies such as ATRA have greatly improved the outlook for AML treatment, its success has been limited to Acute Promyelocytic Leukemia (APL). Therefore, discovering alternative differentiation therapies to treat non-APL patients is a necessity and this discovery of WNK1 inhibitors as potential AML differentiation inducing agents takes us one step forward in that direction.

Our study uncovers a novel WNK1 signaling axis involving the suppression of ERK and C/EBPβ activity. Releasing this suppression through WNK1 inhibition also resulted in release from AML differentiation arrest. Future studies are needed to identify potential additional intermediaries in this signaling axis. It is evident that the effects on ERK are mediated through MEK1/2, but whether WNK1 is directly acting on MEK1/2 or through other protein(s), still needs to be elucidated. This study provides an additional perspective on the role of ERK activation. Traditionally, ERK is thought of as pro-cancerous as it has been identified to activate multiple proliferation associated proteins *(42)*. Additionally, mutations of upstream mediators in this pathway such as BRAF, NRAS, HRAS and KRAS have been well-established as proto-oncogenes *^(43)^*. However, increasing evidence reveals that the role of this signaling axis in cancer is more nuanced than originally believed. ERK hyperactivation has shown the ability to induce apoptosis *(44)* and there has been a recent push to develop ERK agonists to promote these anti-cancerous effects *(45)*. Here we provide insight into how increasing ERK activation can have anti-cancerous effects in the context of AML. We show that there is a threshold at which the level of ERK activation sustains proliferation while also maintaining differentiation arrest and that going above that threshold induces differentiation and counteracts proliferative effects.

This study provides further insight into C/EBPβ biology which also has had seemingly contradictory roles in AML. C/EBPβ expression is strongly induced after treatment with differentiation inducing agents like ATRA *^(29)^* and 1,25D *(35)* and its activity is necessary for ATRA-driven differentiation*^(30)^*. However, other studies have shown that C/EBPβ benefits leukemic growth by coordinating with MYB and p300 to promote transcription of pro-leukemic genes *(46, 47)*. The different effects of C/EBPβ activity may be concentration-dependent with lower levels allowing for pro-leukemic signals and higher levels leading to differentiation-inducing signals. Alternatively, the presence or absence of C/EBPβ co-factors may influence its biological effects. Of note, ATRA treatment decreases expression of MYB which may account for its switch away from pro-leukemic signaling *(48)*. In this study, we show that WNK1 inhibition increases C/EBPβ activity and expression via MEK-ERK to promote differentiation. We observed a robust increase in C/EBPβ expression matching with the aforementioned ATRA and 1,25D studies *^(29, 35)^* that reflect anti-leukemic C/EBPβ functions, further supporting the idea that these effects may depend on C/EBPβ expression levels.

Here we show that AML patients had elevated SPAK expression which correlated with worse overall survival while OSR1 expression was less indicative of disease status and overall survival outcomes. However, we found that both SPAK and OSR1 deficiencies mirrored the deleterious effects of WNK1 inhibition. This could suggest that while higher OSR1 expression does not worsen the disease, its presence is necessary to promote AML growth and progression. These findings are supported by another study which showed that restoring SPAK or OSR1 activity rescued the deleterious effect of WNK1 knockout on AML cell fitness *(23)*. Although SPAK and OSR1 have been thought of as functionally redundant *^(49–51)^*, we observed differential induction of apoptosis and differentiation following their individual knockouts, suggesting functional differences in the context of AML.

Some of the key limitations of this study include the suboptimal inhibitors available for targeting WNK1. While WNK-IN-11 has good specificity for WNK1, its bioavailability is relatively poor. Other available WNK inhibitors, such as WNK-463, also target other WNK-family proteins (WNK2-4). The impact of inhibiting other WNK family may have adverse effects as WNK-463 preclinical safety studies show issues such as ataxia and breathing difficulties *(52)*. Specific WNK1 inhibitors with good pharmacokinetic and pharmacodynamic profiles are needed to further investigate the potential of targeting WNK1 in AML patients. Additionally, our data suggests that WNK1 inhibition may be impacted by the presence of different mutations in AML. We observed cell line specific effects of WNK-IN-11 treatment, especially in apoptosis levels. As these cell lines possess distinct driver mutations, this could indicate that different mutational profiles may impact the efficacy of WNK1 inhibition. However, with the limited number of cell lines used in this study, we are cautious to draw any conclusions about which mutations may make AML cells more or less susceptible to WNK1 inhibition. Thus, future studies testing a broader array of AML cell lines and patient samples are needed to characterize these effects.

Furthermore, as AML heterogeneity is a constant plague to AML treatments, it is imperative that we characterize which subtypes new treatment strategies could be more effective on. Hence, additional studies that further test the potential of WNK1 as a therapeutic target alone or in combination with other therapeutic approaches across multiple AML subtypes are warranted. WNK1 inhibitors are also gaining a lot of attention as a treatment for hypertension *(52, 53)*. As pre-clinical studies are underway for this approach, this may provide an opportunity to repurpose this drug for leukemia treatment.

## MATERIALS AND METHODS

### Mice

NSG mice were purchased from Jackson Laboratories and were housed according to the institutional guidelines at Louis Stokes Cleveland Veterans Affairs Medical center and Case Western Reserve University.

### Ethical approval

All mice were handled in accordance with the National Institutes of Health (NIH) guidelines under Case Western Reserve University protocols (#2013-0134) and Louis Stokes Cleveland Veterans Affairs Medical Center protocol (#2024-0077) approved by the respective Institutional Animal Care and Use Committee. All the experiments were performed in compliance with animal use guidelines (including the ARRIVE guidelines) and the above ethical approval.

### Cell lines

OCI-AML3, MOLM-14, THP-1, HL-60, and MOLM-13 cells were cultured in RPMI-1640 media (Sigma) with 10% FBS, 100 U/mL penicillin/streptomycin, and 292 µg/mL L-glutamine at 37 °C in a humidified 5% CO2 incubator. Routine testing for mycoplasma contamination was performed, and cell line authenticity was confirmed through short tandem repeat (STR) profiling at the Case Western Reserve University genomics core facility.

### Primary HSPC and AML cells

Peripheral blood and bone marrow samples from AML patients and healthy donors were procured through the Case Western Reserve University Hematopoietic Biorepository and Cellular Therapy Core and Pathology lab at Louis Stokes Cleveland VA Medical Center. Ficoll-density purification was used to isolate mononuclear cells (MNCs). For healthy donor bone marrow, cells were further purified using the CD34 MicroBead Kit (Miltenyi) to obtain enriched hematopoietic stem and progenitor cells (HSPCs). Cells from both AML patients and healthy donors were cultured with StemMACS™ HSC Expansion Medium XF (Miltenyi, 130-100-473) at 37 °C and 5% CO2.

### Compounds

WNK-IN-11 (Cayman chemicals, 29676), WNK-463 (MedChem Express, HY-100626), and PD98059 (MedChem Express, HY-12028) were reconstituted in DMSO and aliquoted for storage at -80 C. Inhibitor solutions were prepared fresh in cell culture media for each experiment.

### Lentivirus transduction

For shRNA knockdowns, the following shRNAs were inserted into the pLKO.1 plasmid by Sigma Aldrich and delivered to us as glycerol stocks: WNK1 – CCGCGATCTTAAATGTGACAA (TRCN0000000919), STK39 (SPAK) – CGGTCAGATTCACAGGGATTT (TRCN0000001006), OXSR1 (OSR1) - GCGTATCTCTGTTGCTTCTAT (TRCN0000001586). HEK-293 cells were co-transfected with shRNA-containing plasmids (5 µg), psPAX2 (3.3 µg), and pMD2.G (1.8 µg) in a 100 mm tissue culture plate. Lentiviral particles were harvested from culture supernatant 48 hours post-transfection and concentrated by ultracentrifugation. 1 x 10^6^ cells were spin-infected with the lentivirus concentrate by centrifuging at 2500 rpm for 90 minutes at 30 °C in the presence of 7.5 µg/mL polybrene (Sigma, TR1003G). After spin-infection, the media was replaced. For studies on cell cycle, apoptosis, and differentiation, the cells were analyzed 4 days after infection.

### Flow cytometry

#### Cell Cycle Analysis

MOLM-14, OCI-AML3, MOLM-13 (1.5 x 10^4^) and THP-1 (3 x 10^4^) cells were cultured with WNK-IN-11 (10 µM) or vehicle control for 3 days. Cells were permeabilized with 70% ethanol for 30 minutes at 4°C then washed twice with PBS. 20 µg/mL Propidium iodide (PI) (BioLegend, 421301) and 100 µg/mL RNase (Sigma, R6148) were added to the cell suspension. Samples were incubated at room temperature in the dark for 15 minutes.

#### Annexin V/PI staining

MOLM-14, OCI-AML3, MOLM-13 (7.5 x 10^3^) and THP-1 (1.5 x 10^4^) cells were cultured with WNK-IN-11 (10 µM) or vehicle control for 4 days. Staining solution was prepared by combining Annexin V APC (BioLegend, 640941) to Annexin V binding buffer (BioLegend, 422201). Cells were resuspended in staining solution. Cells were additionally stained with 20 ug/ml PI and incubated in the dark at room temperature for 30 minutes.

#### Differentiation marker quantification

MOLM-14, OCI-AML3, MOLM-13, and THP-1 (3 x 10^5^) cells were cultured with WNK-IN-11 (10 µM) or DMSO control for 72 hours. The cells were pre-incubated with TruStain Human FcX Blocking Buffer (BioLegend, 422302) prior to staining with anti-CD11b (BioLegend, 101212) and anti-CD14 antibodies (BioLegend, 367104). Dead cells were excluded using Zombie NIR Fixable dye.

#### Analysis

Samples were analyzed with a CytoFLEX flow cytometer (Beckman Coulter) and FlowJo software v10.8.1.

### Wright-Giemsa staining

3 x 10^5^ cells were cultured with WNK-IN-11 (10 µM) or DMSO control for 72 hours. 5 x 10^4^ cells were transferred to slides using the Shandon Cytospin 3 centrifuge (800 rpm for 3 minutes). Slides were stained with Wright-Giemsa by the University Hospitals Hematopathology and Flow Cytometry Core. Slides were then visualized on an Olympus IX73 microscope.

### Cell proliferation assay

MOLM-14, OCI-AML3, MOLM-13 (7.5 x 10^3^) and THP-1 (1.5 × 10^4^) cells were cultured with WNK-IN-11 (10 µM) or DMSO control for 4 days. Each day, cells were collected and mixed with 0.4% Trypan blue at a 1:1 ratio. Cell counts were collected using a Countess 3 Automated Cell Counter (ThermoFisher).

### OCI-AML3 mouse xenograft

NSG mice were injected with 1 x 10^6^ MOLM-14 or OCI-AML3 cells into the lateral tail vein and allowed to engraft. Female NSG mice were used for xenograft studies because of their higher engraftment efficiency compared to males *(54)* and assigned randomly to treatment or control groups. After 7 days, engraftment was confirmed via bioluminescent imaging and mice were treated with 2 cycles of 5-day treatment and 2 days rest with WNK-463 (1.5 mg/kg) or vehicle control. Mice were imaged at day 7, 14, and 21 post-injection and were sacrificed at the conclusion of the study.

### Quantitative PCR

RNA was isolated using the EZ10 DNAaway RNA miniprep kit (BioBasic). 1 µg of total RNA was converted to cDNA using the High Capacity cDNA Reverse Transcription Kit (Applied Biosystems). Quantitative real-time PCR was performed using PowerUP SYBR Green Master Mix (Applied Biosystems) following manufacturer’s protocol. Gene expression was normalized to the RPL32 housekeeping gene and was quantified using the 2-ΔΔCt method *^(55)^*. Primer sequences are included in Table S3.

### Western blotting

Cell lysates were extracted using Triton Lysis Buffer (1% Triton-X100, 20 mM HEPES [pH 7.6], 0.1% SDS, 0.5% Sodium deoxycholate, 150 mM NaCl, 1 mM EDTA) supplemented with protease inhibitor cocktail (ThermoFisher, Cat# A32955). Lysates were resolved through 7%, 9%, or 15% SDS-PAGE gels, transferred to a nitrocellulose membrane, and probed using the following antibodies: WNK1 (Cell Signaling, 4979S), SPAK (Santa Cruz, sc-517361), OSR1 (Santa Cruz, sc-271707), phospho SPAK/OSR1 (Sigma, 07-2273), C/EBPβ (Cell Signaling, 43095S), phospho C/EBPβ (Cell Signaling, 3084S), Vinculin (Santa Cruz, sc-73614), PARP (Santa Cruz, sc-7150), Caspase 3 (Cell Signaling, 9662S), Cleaved caspase 3 - Cell Signaling, 9661S, HDAC2 (Santa Cruz, sc-6296), β-Tubulin (Santa Cruz, sc-9104), ERK1/2 (Cell Signaling, 9102S), phospho ERK1/2 (Cell Signaling, 4370S) followed by anti-mouse (ThermoFisher, 31430) and anti-rabbit (ThermoFisher, 31460) secondary antibodies conjugated with horse radish peroxidase (HRP) and then visualized using enhanced chemiluminescent substrate (GenDepot) x-ray film imaging. For western blotting assessing increases in C/EBPβ and ERK1/2 phosphorylation, cells were serum starved 16 hours prior to inhibitor treatment. After the indicated treatment periods, lysates were collected and subject to western blot analysis as described above.

### Oligonucleotide pulldown assay

Oligonucleotide pulldown assay was performed as described previously *(56)*. Cytoplasmic and nuclear-enriched fractions were prepared by lysing in hypotonic cytoplasmic lysis buffer (10 mM HEPES pH 7.6, 10 mM KCl, 0.1 mM EDTA, 0.1 mM EGTA) supplemented with protease inhibitor cocktail for 15 min on ice. NP40 (0.625%) was added, and lysates were centrifuged at 12,000 g for 30 seconds. Supernatants were collected as the cytoplasmic-enriched fractions. The pellets were washed once with hypotonic cytoplasmic lysis buffer. Nuclear pellets were then lysed with hypertonic nuclear lysis buffer (20 mM HEPES pH 7.6, 400 mM NaCl, 1 mM EDTA, 1 mM EGTA) supplemented with protease inhibitor cocktail for 30 min on ice. Lysates were centrifuged at 12,000 g for 10 min at 4 °C and supernatants were collected as the nuclear-enriched fractions. NaCl concentration was adjusted to 150 mM and 50 µg of nuclear lysate was incubated with 1 µg of biotinylated oligonucleotides, 6 µg of sheared salmon sperm DNA to block nonspecific binding, and Neutravidin agarose beads (ThermoFisher) for 16 hours at 4 °C with gentle rotation. Beads were washed 3 times with nuclear washing buffer (20 mM HEPES pH 7.6, 150 mM NaCl, 1 mM EDTA, 1 mM EGTA). Samples were boiled at 95 °C for 5 mins and centrifuged at 10,000 x g for 5 minutes. Supernatant was then resolved through a 9% SDS-PAGE gel and analyzed by western blotting.

### Overall survival

Overall survival was analyzed using SurvivalGenie *(24)*. Peripheral blood samples from the TARGET cohort were stratified based on WNK1, SPAK, or OSR1 expression with high expressors defined as those top 20th percentile and low expressors as those in the bottom 20th percentile.

### RNA sequencing

OCI-AML3 cells (7.5 x 10^5^) were cultured with WNK-IN-11 (10 µM) or DMSO Ctrl for 12 hours and RNA was extracted using the RNAeasy Mini Kit (Qiagen). Library preparation and whole transcriptome sequencing was performed by Novogene using 3 replicates from each group. Differentially expressed genes were defined as those with a log2FoldChange > 1 and an adjusted p-value < 0.05. KEGG enrichment analysis were done using the NovoMagic cloud platform (https://magic.novogene.com/customer/main#/login). GSEA was performed using the Broad Institute web platform *(57, 58)* with a pre-ranked list from the gene expression data based on log2FoldChange.

### Phosphoproteomic mass spectrometry

OCI-AML3 cells were treated with DMSO or WNK-IN-11 (10 µM) for 3 hours. Triplicate lysates per condition were collected using 2% SDS solution supplemented with protease inhibitor cocktail (Sigma, P2714) and PhosphoSTOP (Roche, 04906837001). Lysates were then processed by following the filter assisted sample preparation (FASP) cleaning procedure as previously described *(59)* and were normalized to 500 µg. Peptides were generated by digestion with dual LysC (Wako, 125-05061) and trypsin (ThermoFisher, 90057). The digested samples were desalted using C18 cartridges (Oasis C18 1cc Waters) and subjected to the High Select™ Phosphopeptide enrichment kit (ThermoFisher, A32993). Mass spectrometry data was collected using LC-MS/MS via a UPLC Vanquish chromatography system (ThermoFisherScientific) coupled with an Orbitrap 480 Exploris mass spectrometer (ThermoFisherScientific). Mass spectrometry data was processed using PEAKS v10.6 software (Bioinformatics Solutions) and peptide identification was performed using the UNIPROT database (human UP000005640). Label-free quantification was performed using default settings, which included mass error tolerance of 10ppm, retention time shift tolerance of 5.0 min, 0.6 Da for fragment ions, and TIC normalization. Differentially phosphorylated phosphopeptides were classified as those with a p-value < 0.05 and Fold change > 2 (upregulated) or < 0.5 (downregulated) and visualized with the ggplot2 package in R. Kinase-substrate enrichment analysis was done using the KSEA App online tool (https://casecpb.shinyapps.io/ksea/) *(31–34)*.

### Colony Formation Assay

Mononuclear cells from AML bone marrow were plated at 2 x 10^4^ and 4 x 10^4^ cells/mL in MethoCult media (Stemcell Technologies, H4435) according to manufacturer’s instructions. On day 14, colonies from plates were counted blind and pictures were acquired using a Cytation 5 microscope (BioTek).

### Chloride uptake

OCI-AML3 cells (2 x 10^6^) were resuspended in a hypotonic medium composed of Hank’s Balanced Salt Solution (HBSS) and Sterile-filtered water combined at a 1:1 ratio. Cells were then loaded with 5 mM MQAE for 30 minutes at 37°C. Cells were then spun down and resuspended in HBSS with 0, 5, 10, or 50 µM bumetanide and incubated for another 30 minutes at 37°C. Chloride influx was assessed by the quenching of MQAE fluorescence (excitation/emission, 350/460) measured in each well on a Spectramax i3x Multi Mode microplate reader (Molecular Devices). The chloride uptake index was calculated by taking the inverse of the MQAE fluorescent units.

### Statistical analyses

Data were analyzed with GraphPad Prism (v10.2.2). A Student’s t-test was used for comparisons between two groups. One-way ANOVA with Dunnet’s multiple comparison test (when comparing each treatment group to the control) or Tukey’s multiple comparison test (when comparing multiple treatment groups to each other) were used to compare more than two groups. Two-way ANOVA with Dunnet’s multiple comparison test (when comparing each treatment group to the control) or Šídák’s multiple comparisons test (when comparing two treatment groups) were used for pairwise comparisons. Dose-response curves were calculated by plotting DMSO control normalized cell viability fitted with a non-linear regression model. GraphPad was used to calculate the IC_50_ values.

AML samples from TCGA, BeatAML, TARGET, and St. Jude Children’s Research Hospital’s St. Jude Cloud (SJC-DS-1013 and SJC-DS-1009) and healthy whole blood from GTEx and HSPCs from GSE198919 were compiled as described previously *(60)* and imported into R. (v4.5.1) Genes not found across all datasets were removed. Counts were normalized using voom from limma v3.54.2 *(61)*. Differential expression was calculated using eBayes and topTable *(62, 63)*.

For WNK-IN-11 RNA sequencing analysis, the edgeR and DESeq2 packages were used in R. Count data were fitted to a negative binomial generalized linear model. Differential expression was assessed using the Wald test and p-values were adjusted using the Benjamini-Hochberg correction. WNK-IN-11 phosphoproteomics was assessed by calculating the significance score and p-value in PEAKS v10.6 software (Bioinformatics Solutions) using an FDR threshold of 2%.

## Acknowledgments

We thank the CWRU biorepository core and VA pathology lab for providing AML patient cells. We thank the CWRU Proteomics and Small Molecule MS core (NIH, S10OD026882) for performing phosphoproteomic mass spectrometry. We thank CWRU Imaging Research core for BLI assistance, Drs. Derek Abbot and Reshmi Parameswaran for kindly providing MOLM-14 and OCI-AML3 cells, and Drs. David Wald, Alex Huang, George Dubyak, and John Letterio for insightful discussions.

## Funding

U.S. Department of Veteran Affairs VA BLRD I01 BX005941 (PR), National Institutes of Health (NIH)/National Cancer Institute (NCI) grant R21CA246194 (PR), NIH/National Institute of Diabetes and Digestive and Kidney Diseases (NIDDK) R01DK128463 (PR), Breakthrough T1D (Formerly (Juvenile Diabetes Research Foundation, JDRF) 3-SRA-2022-1193-S-B (PR), JDRF 3-SRA-2024-1552-S-B (PR), NIH/NIAID T32-AI089474 (JDC).

## Author contributions

Conceptualization: JDC, PR

Methodology: JDC, PR

Investigation: JDC, EMK, PR

Visualization: JDC, EMK, PR

Funding acquisition: JDC, PR

Project administration: JDC, PR

Supervision: JDC, PR

Writing – original draft: JDC, EMK, PR

Writing – review & editing: JDC, EMK, PR

## Competing interests

Authors declare that they have no competing interests.

## Data and materials availability

All the data and methods necessary to reproduce this study are included in the manuscript and supplementary materials. Reagent and protocol requests as well as additional details on proteomics mass spectrometry will be readily fulfilled following the materials transfer policies of Case Western Reserve University (CWRU). WNK-IN-11 RNA sequencing data has been deposited in the Gene Expression Omnibus under accession code GSE318062. The WNK-IN-11 mass spectrometry data and KSEA analysis have been included as supplementary files (Data files S1-2). Publicly available datasets used in this study: GTEx (via www.gtexportal.org); TCGA, BeatAML, and TARGET (via GDC commons); St. Jude Cloud (via platform.stjude.cloud/data, accession numbers SJC-DS-1013 and SJC-DS-1009) and GSE198919, GSE116256, and GSE126068 via GEO). For inquiries, please contact Parameswaran Ramakrishnan at.

## Notes

https://www.ncbi.nlm.nih.gov/geo/query/acc.cgi?acc=GSE318062

### Competing Interest Statement

The authors have declared no competing interest.

