## Supplemental figures for "Targeting WNK1 Releases Differentiation Block in Acute Myeloid Leukemia"

### Slide 1
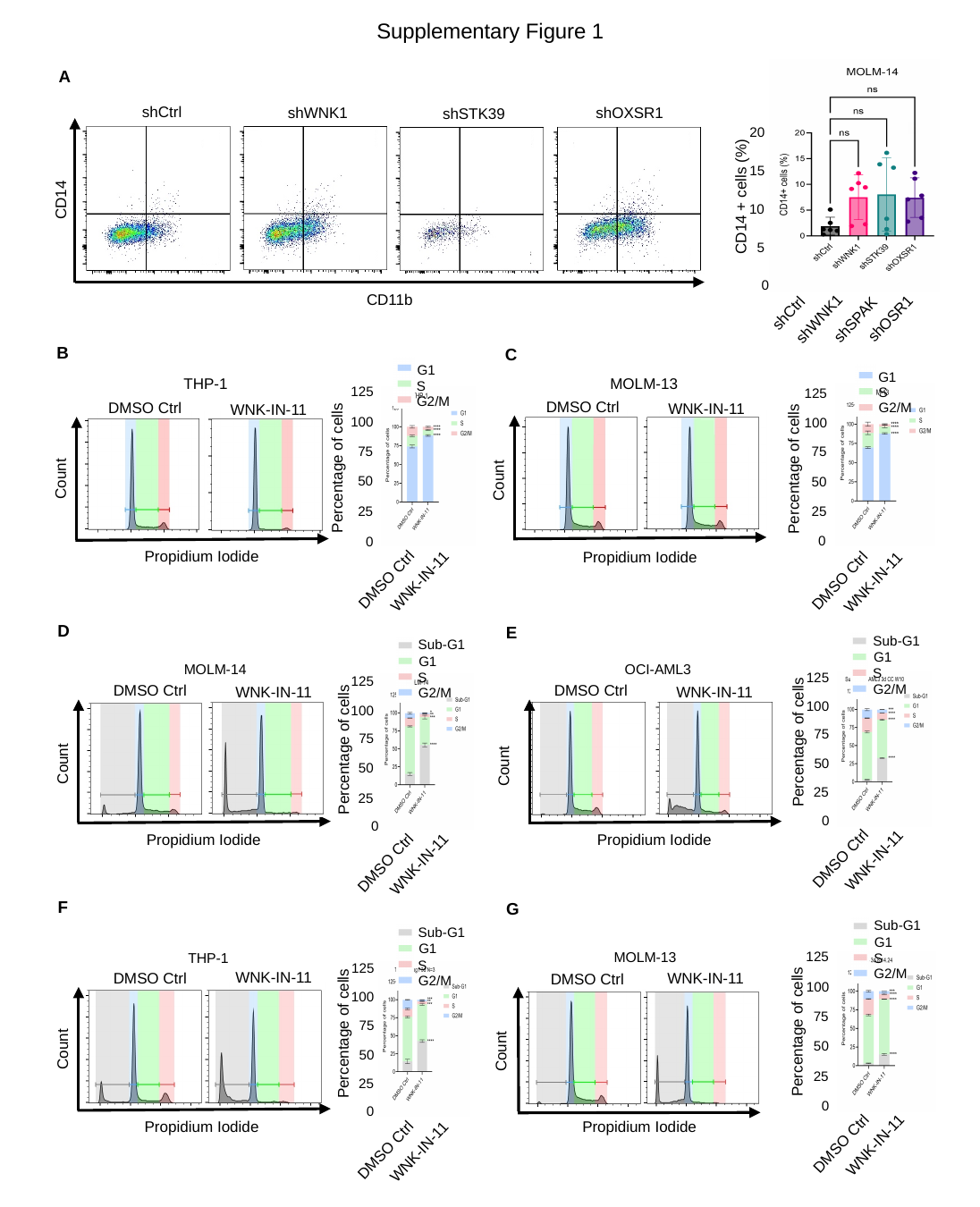

#### Supplementary Figure 1
A
shCtrl
shWNK1
shOXSR1
shSTK39
B
20
15
CD14 + cells (%)
CD14
10
5
0
CD11b
shCtrl
shOSR1
shWNK1
shSPAK
B
C
G1
G1
THP-1
MOLM-13
S
125
S
125
G2/M
DMSO Ctrl
DMSO Ctrl
WNK-IN-11
G2/M
WNK-IN-11
100
100
75
75
Percentage of cells
Percentage of cells
Count
Count
50
50
25
25
0
0
Propidium Iodide
Propidium Iodide
DMSO Ctrl
DMSO Ctrl
WNK-IN-11
WNK-IN-11
D
E
Sub-G1
Sub-G1
G1
G1
OCI-AML3
MOLM-14
S
125
S
125
G2/M
DMSO Ctrl
DMSO Ctrl
WNK-IN-11
WNK-IN-11
G2/M
100
100
75
75
Percentage of cells
Percentage of cells
Count
50
Count
50
25
25
0
0
Propidium Iodide
Propidium Iodide
DMSO Ctrl
WNK-IN-11
DMSO Ctrl
WNK-IN-11
F
G
Sub-G1
Sub-G1
G1
G1
125
MOLM-13
THP-1
S
S
125
G2/M
WNK-IN-11
DMSO Ctrl
WNK-IN-11
DMSO Ctrl
G2/M
100
100
75
75
Percentage of cells
Percentage of cells
50
Count
Count
50
25
25
0
0
Propidium Iodide
Propidium Iodide
DMSO Ctrl
WNK-IN-11
DMSO Ctrl
WNK-IN-11

### Slide 2
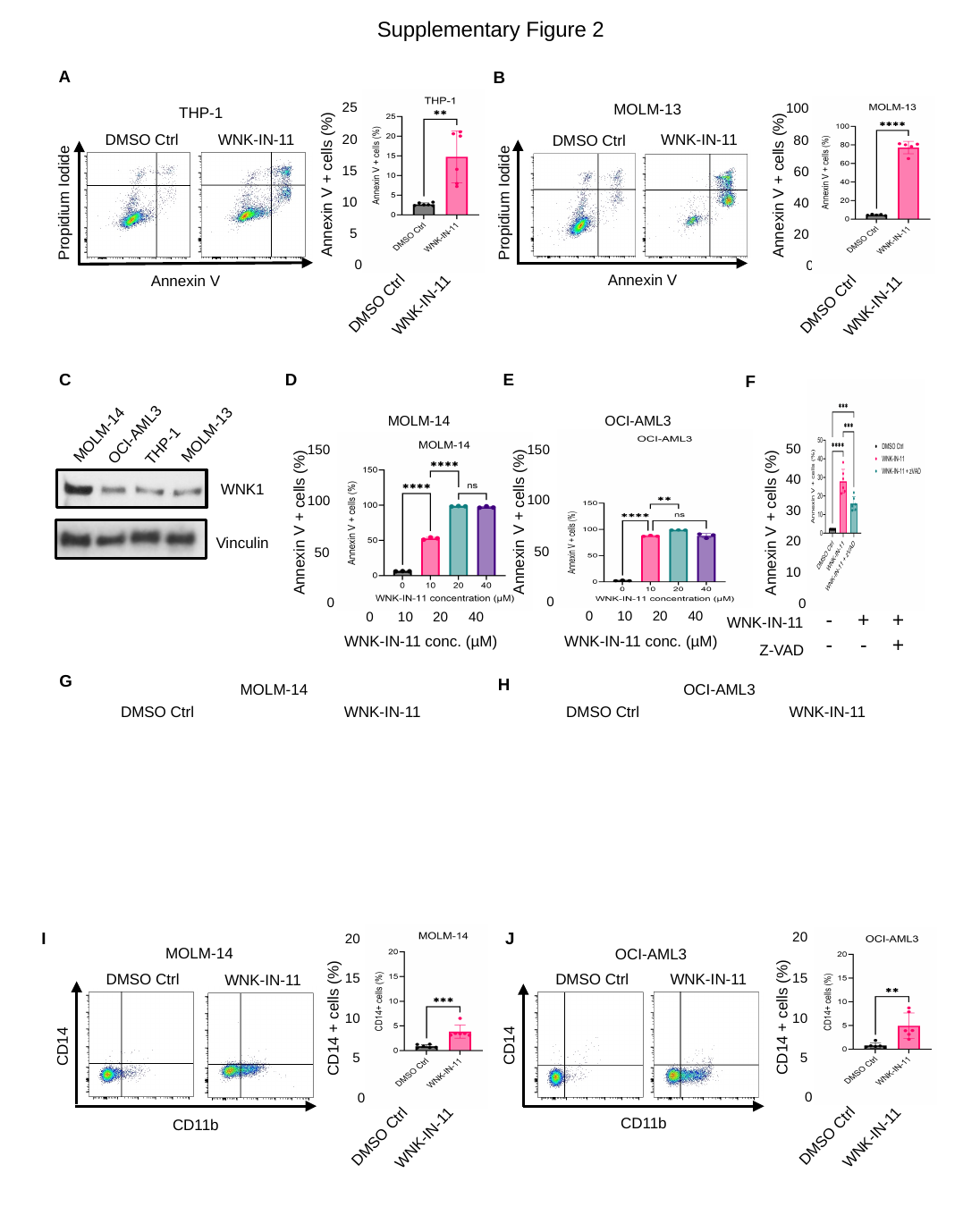

#### Supplementary Figure 2
A
B
25
100
MOLM-13
THP-1
WNK-IN-11
DMSO Ctrl
20
WNK-IN-11
DMSO Ctrl
80
15
60
Annexin V + cells (%)
Annexin V + cells (%)
Propidium Iodide
Propidium Iodide
10
40
5
20
0
0
Annexin V
Annexin V
DMSO Ctrl
DMSO Ctrl
WNK-IN-11
WNK-IN-11
E
D
C
F
MOLM-14
OCI-AML3
OCI-AML3
MOLM-14
MOLM-13
THP-1
50
150
150
40
WNK1
100
100
30
Annexin V + cells (%)
Annexin V + cells (%)
Annexin V + cells (%)
20
Vinculin
50
50
10
0
0
0
20
40
0
10
20
40
0
10
| - | + | + |
| --- | --- | --- |
WNK-IN-11
WNK-IN-11 conc. (µM)
WNK-IN-11 conc. (µM)
| - | - | + |
| --- | --- | --- |
Z-VAD
G
H
OCI-AML3
MOLM-14
DMSO Ctrl
WNK-IN-11
DMSO Ctrl
WNK-IN-11
I
J
20
20
MOLM-14
OCI-AML3
15
DMSO Ctrl
15
DMSO Ctrl
WNK-IN-11
WNK-IN-11
CD14 + cells (%)
CD14 + cells (%)
10
10
CD14
CD14
5
5
0
0
CD11b
CD11b
DMSO Ctrl
DMSO Ctrl
WNK-IN-11
WNK-IN-11

### Slide 3
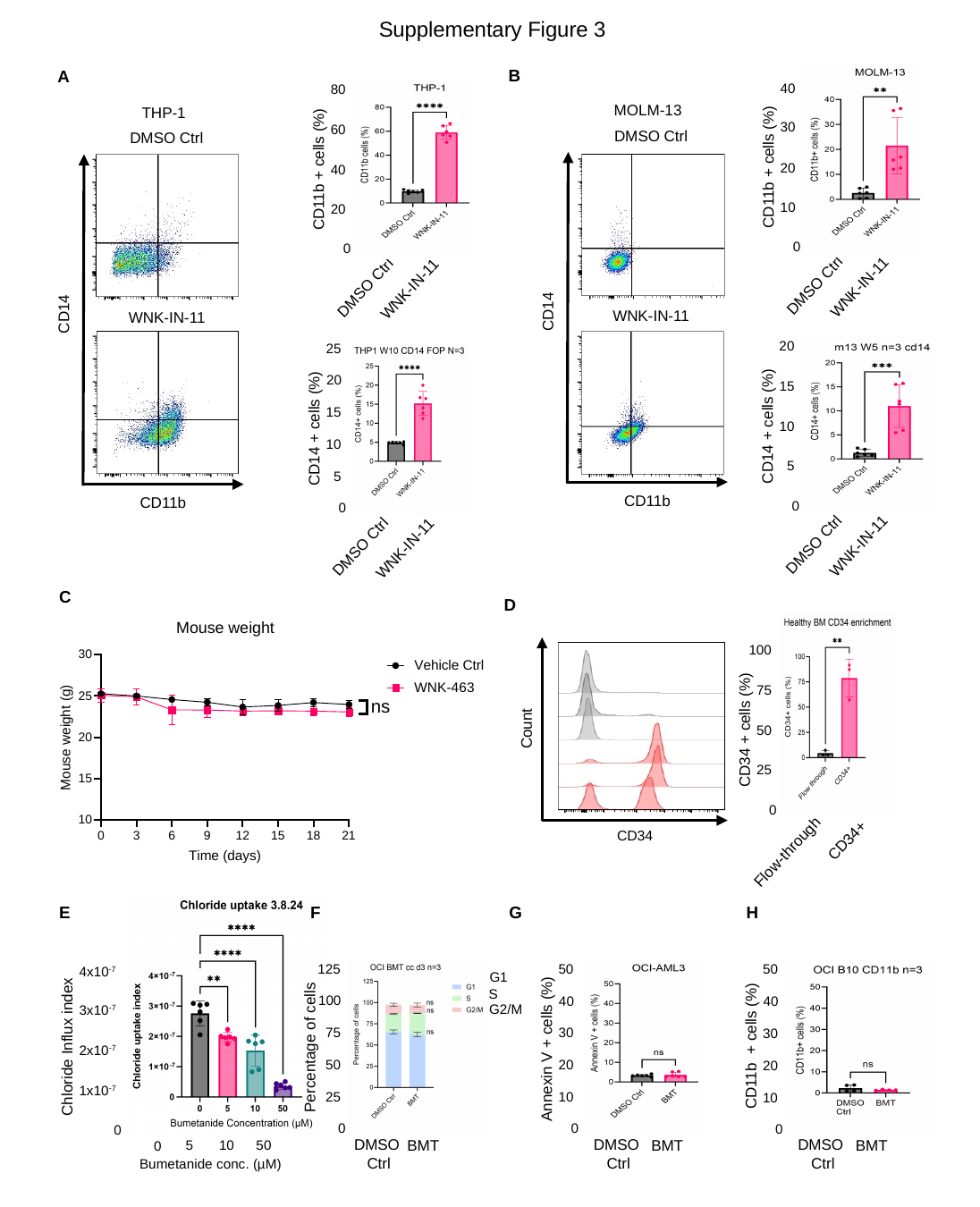

#### Supplementary Figure 3
B
A
40
80
MOLM-13
THP-1
30
60
DMSO Ctrl
DMSO Ctrl
CD11b + cells (%)
20
CD11b + cells (%)
40
10
20
0
0
DMSO Ctrl
WNK-IN-11
DMSO Ctrl
WNK-IN-11
CD14
CD14
WNK-IN-11
WNK-IN-11
20
25
20
15
15
CD14 + cells (%)
10
CD14 + cells (%)
10
5
5
CD11b
CD11b
0
0
DMSO Ctrl
WNK-IN-11
DMSO Ctrl
WNK-IN-11
C
D
100
75
Count
CD34 + cells (%)
50
25
0
CD34
CD34+
Flow-through
E
F
H
G
125
50
50
4x10^-7^
G1
S
100
40
40
G2/M
3x10^-7^
75
30
30
CD11b + cells (%)
Chloride Influx index
Percentage of cells
Annexin V + cells (%)
2x10^-7^
50
20
20
1x10^-7^
25
10
10
0
0
0
0
DMSO
Ctrl
DMSO
Ctrl
DMSO
Ctrl
BMT
BMT
BMT
5
50
10
0
Bumetanide conc. (µM)

### Slide 4
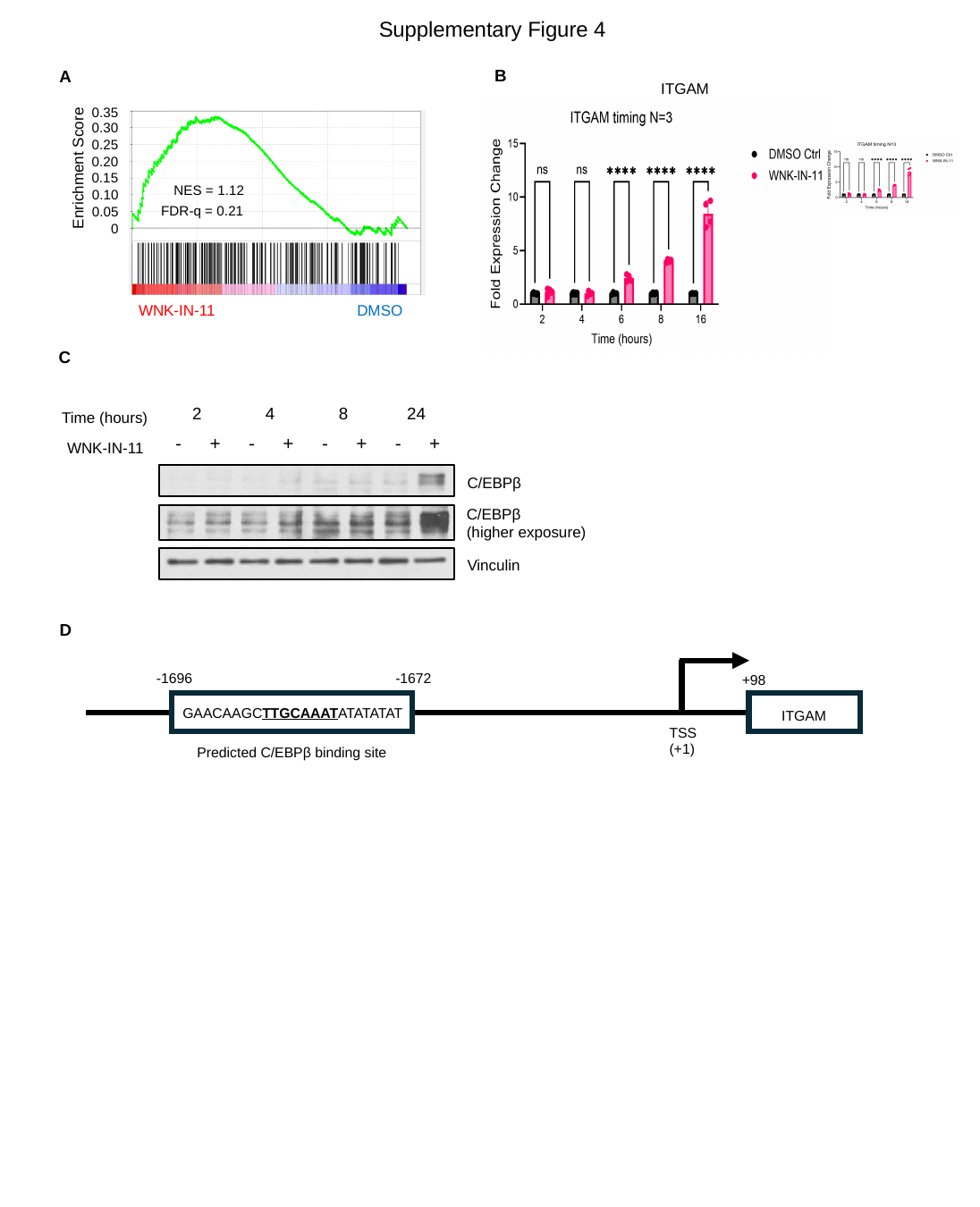

#### Supplementary Figure 4
B
A
ITGAM
0.35
0.30
0.25
0.20
Enrichment Score
0.15
NES = 1.12
0.10
FDR-q = 0.21
0.05
0
WNK-IN-11
DMSO
C
| 2 | 4 | 8 | 24 |
| --- | --- | --- | --- |
Time (hours)
| - | + | - | + | - | + | - | + |
| --- | --- | --- | --- | --- | --- | --- | --- |
WNK-IN-11
C/EBPβ
C/EBPβ
(higher exposure)
Vinculin
D
-1696
-1672
+98
GAACAAGCTTGCAAATATATATAT
ITGAM
TSS
(+1)
Predicted C/EBPβ binding site
