## Supplemental figure legends and tables for "Targeting WNK1 Releases Differentiation Block in Acute Myeloid Leukemia"

Supplementary Materials for **Targeting WNK1 Releases Differentiation Block in Acute Myeloid Leukemia**

**This document includes:**

Figs. S1-4

Tables S1-3

**Other Supplementary Material for this manuscript includes the following:**

Data file S1-3

Fig S1. A) (Left) Scatter plots showing CD11b and CD14 surface expression on MOLM-14 cells 4 days after lentiviral shRNA transduction. (Right) Frequency of CD14+ MOLM-14 cells assessed 4 days after lentiviral shRNA transduction. B-C) THP-1 (B) and MOLM-13 (C) cells were treated with WNK-IN-11 (10 µM) or DMSO control for 3 days prior to cell cycle analysis. (Left) Histogram showing cellular DNA content by propidium iodide staining. (Right) Percentages of cells in each phase of cell cycle. D-G) Cell cycle analysis 3 days after WNK-IN-11 not excluding dead cells for MOLM-14 (D), OCI-AML3 (E), THP-1 (F), and MOLM-13 (G) cells. (Left) Histogram showing cellular DNA content by propidium iodide staining. (Right) Percentages of cells in each phase of cell cycle. Statistical analysis was done using a one-way ANOVA with Dunnet’s multiple comparisons test (A) or a two-way ANOVA with Šídák's multiple comparisons test (B-G). **, P < 0.01; ****, P < 0.0001; ns, not significant.

Fig. S2. A-B) Annexin V positivity assessed on THP-1 (A) and MOLM-13 (B) cells after 4 days of WNK-IN-11 (10 µM) or DMSO Control treatment. (Left) Annexin V and PI staining for apoptosis, (Right) Percentage of Annexin V positive cells. C) Western blot showing levels of WNK1 in MOLM-14, OCI-AML3, THP-1, and MOLM-13 cells. D-E) Annexin V positivity assessed with increasing concentrations of WNK-IN-11 treatment on MOLM-14 (D) and OCI-AML3 (E) cells. E) Annexin V positivity on OCI-AML3 cells after 3 days of treatment with WNK-IN-11 (10 µM) or a combination of WNK-IN-11 and pan-caspase inhibitor, Z-VAD (20 µM). G-H) Wright-Giemsa stain showing morphological changes of MOLM-14 (G) and OCI-AML3 (H) 3 days after treatment with WNK-IN-11 (10 µM) or DMSO control (magnification: 60x). I-J) CD11b and CD14 surface expression 3 days after WNK-IN-11 treatment on MOLM-14 (I) and OCI-AML3 (J) cells. (Left) Scatter plots showing CD11b and CD14 positivity. (Right) Percentage of CD14+ cells. CD14 expression is represented graphically on the right. Statistical analysis was done using a Student’s t-test (A-B, I-J) one-way ANOVA with Tukey's multiple comparisons test (D-F). **, P < 0.01; ***, P < 0.001; ****, P < 0.0001.

Fig. S3. CD11b and CD14 surface expression 3 days after WNK-IN-11 treatment on THP-1 (A) and MOLM-13 (B) cells. (Left) Scatter plots showing CD11b and CD14 positivity. (Right) Percentage of CD11b+ and CD14+ cells. C) Mouse weight monitored over 3 weeks of treatment with WNK-463 (1.5 mg/kg) or vehicle control. D) Validation of CD34 enrichment after magnetic isolation. (Left) Histogram overlay of CD34 expression in the flow through (grey) or enriched fraction (red). (Right) Percentage of CD34+ cells in each fraction. E) Inhibition of chloride influx after 30 minutes of Bumetanide treatment quantified by MQAE fluorescence. F) Cell cycle analysis in OCI-AML3 cells 3 days after Bumetanide (10 µM) or DMSO Control treatment. G) Apoptosis markers assessed on OCI-AML3 cells after 4 days of Bumetanide (10 µM) or DMSO Control treatment. H) CD11b surface expression on OCI-AML3 cells 3 days after Bumetanide (10 µM) treatment. Statistical analysis was done using a Student’s t-test (A-D, G-H), one-way ANOVA with Dunnet’s multiple comparisons test (E), or two-way ANOVA with Šídák's multiple comparisons test. **, P < 0.01; ***, P < 0.001; ****, P < 0.0001; ns, not significant.

Fig. S4. A) Gene set enrichment analysis of C/EBPβ target genes in WNK-IN-11 treatment compared to DMSO Ctrl. B) ITGAM gene expression levels 2-16 hours after WNK-IN-11 treatment. C) C/EBPβ expression kinetics following WNK-IN-11 treatment (10 µM) from 2 to 24 hours. D) Oligonucleotide pull-down showing C/EBPβ DNA binding capacity to its consensus sequence (TTGCGCAA) or mutated sequence (CCGAGCAG) in OCI-AML3 cells 2 hours after treatment with WNK-IN-11 (10 µM) or DMSO control. The sequences of the wild-type and mutant oligos are included with the C/EBPβ binding motif in bold and the mutations introduced in red. E) Diagram showing the predicted C/EBPβ binding site used to make the ITGAM promoter oligonucleotide.

**Table S1**

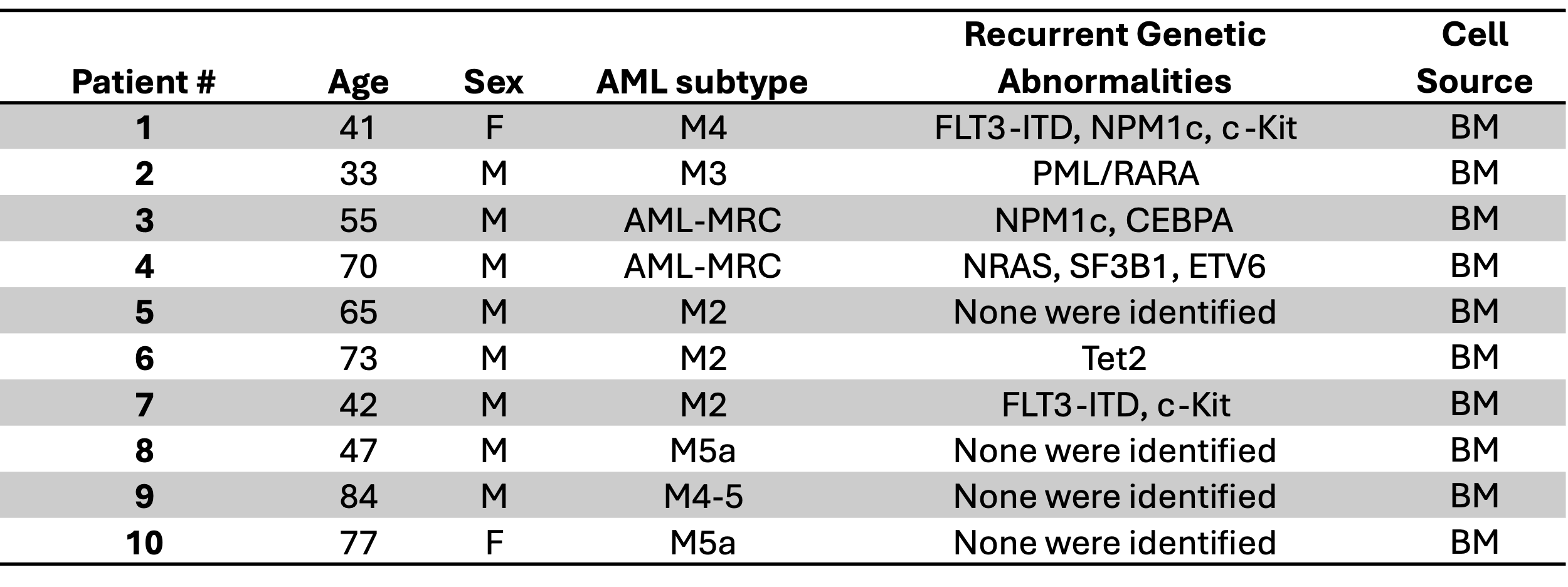

Table S1. Patient demographics for AML samples.

Available information on AML patient demographics and genetic abnormalities. AML-MRC, AML with myelodysplasia related changes; BM, Bone marrow.

**Table S2**

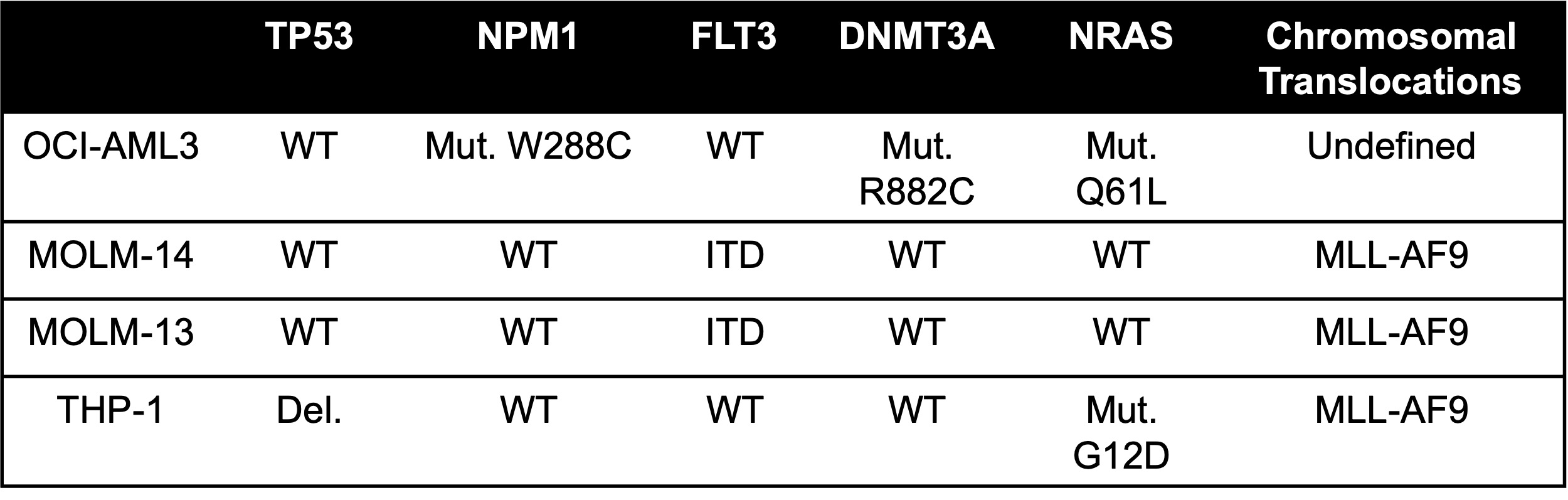

Table S2. AML cell line genetic characterization.

Identification of mutations and chromosomal translocations present in each of the cell lines used in this study.

**Table S3**

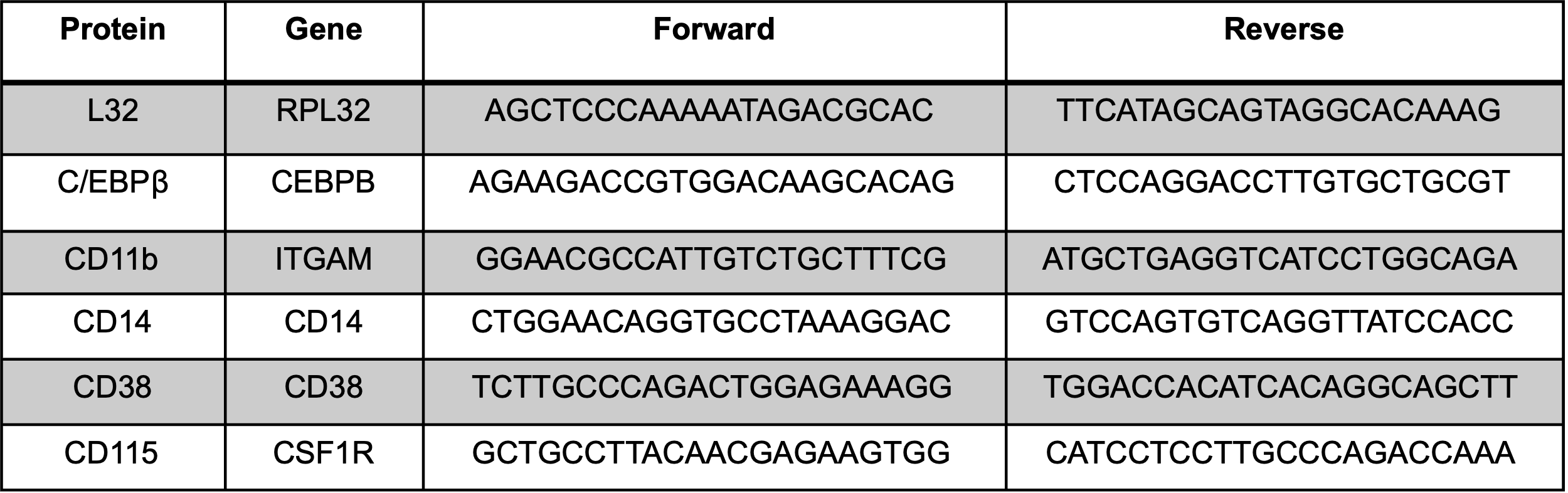

Table S3. Sequences of qPCR primers used.
